# Efficient NK cell transduction with VSV-G-pseudotyped lentiviral vectors

**DOI:** 10.64898/2026.03.11.710988

**Authors:** Emmi Järvelä, Jan Koski, Farhana Jahan, Antti Tuhkala, Mira Saari, Manar Elmadani, Kari Salokas, Laurens Veltman, Lotta Andersson, Hekim Can, Marie Nyman, Seppo Ylä-Herttuala, Markku Varjosalo, Diana Schenkwein, Henrik Paavilainen, Kim Vettenranta, Matti Korhonen, Helka Göös

## Abstract

The need for safe, allogeneic cell therapies for cancer is driving a growing interest in CAR-NK-based therapies, which, unlike CAR-T cell therapies, offer the potential for off-the-shelf administration. Lentiviruses pseudotyped with vesicular stomatitis virus glycoprotein G (VSV-G) are commonly used for genetic modification of cell therapy products. Their use in NK cells, however, is limited by low transduction efficiency. This study explores the complexities of NK cell transduction using lentiviral vectors pseudotyped with VSV-G. We demonstrate that efficient transduction depends on multiple factors such as NK cell activation, construct design, lentivirus pseudotype selection, and the use of transduction enhancers. By optimizing these elements, we achieved effective transduction, facilitating the use of VSV-G-pseudotyped LVs for therapeutic NK cell production. Our optimized workflow comprises NK cell activation with interleukins, followed by transduction with a NK cell-specific CAR construct using VSV-G-pseudotyped LVs in the presence of BX795 and Retronectin, resulting in excellent transduction efficiency without compromising NK cell phenotype or growth. This allows for the use of a widely used gene transfer vector with an excellent safety record for producing therapeutic NK cell products.

## Introduction

T cells equipped with chimeric antigen receptors (CAR-T cells) have yielded impressive results in the treatment of haematological malignancies. They are, however, associated with severe toxicities such as cytokine release syndrome (CRS) and immune effector cell– associated neurotoxicity syndrome (ICANS), and the autologous nature of CAR-T therapies poses logistical and financial challenges. Conversely, allogeneic CAR-NK cells may be associated with fewer toxicities and be manufactured as off-the-shelf products.^1^ The FDA and EMA have also issued warnings about the risk of secondary malignancies following CAR-T cell therapies, highlighting the need for safer options.^2,3^ In contrast to T cells, however, CAR transfer to NK cells is highly challenging.^4^ Current approaches to combat this challenge include retro- or lentiviral transduction, electroporation and oligomers.^5^

HIV-derived lentiviral vectors (LVs) pseudotyped with vesicular stomatitis virus glycoprotein G (VSV-G) are widely used for the genetic modification of therapeutic cells ^6,7^ and their safety and stability have been documented in clinical trials.^6,8^ VSV-G-pseudotyped LVs enter their target cells via the low-density lipoprotein receptor (LDLR).^9^ Freshly isolated NK cells express LDLR poorly, but upregulate its expression upon activation.^10^ Even so, LDLR expression remains low in activated NK cells compared to T cells.^11^ Recently, Gong et al. reported that rosuvastatin upregulates the expression of LDLR on NK cells and can be harnessed to improve the transduction of NK cells with VSV-G-pseudotyped LV.^10^ Adding IL-2 and rosuvastatin to NK cells prior to transduction resulted in a transduction efficiency of 33,3% using VSV-G LVs carrying green fluorescent protein (GFP) as the transgene. However, a study by Allan et al. achieved a similar level of transduction using only IL-2 pre-activation.^12^

In addition to pre-activation and the use of rosuvastatin, a variety of methods can enhance the transduction of NK cells with VSV-G-pseudotyped LVs. Notably, the inhibition of the TBK1/IKKε antiviral signalling pathway using compounds such as amlexanox, MRT67307, or BX795 significantly enhances NK cell transduction. Chockley et al. demonstrated that combining these inhibitors with IL-2 pre-activation can yield a YFP expression of up to 47% in NK cells.^13^ Furthermore, agents like Retronectin and Vectofusin-1 physically concentrate the vector on the NK cell surface.^14,15^ Retronectin is a recombinant fragment of human fibronectin and contains the binding domains for both the vector and NK cell, thereby facilitating closer virus-cell interaction and enhancing transduction efficiency.^14^ Similarly, Vectofusin-1 aggregates the vector on the NK cell through nanofiber formation. It also offers the advantage of not requiring attachment to the cell culture dish surface.^15^

To overcome the challenges in efficient NK cell transduction, this study aims to investigate various methods and their combinations to boost VSV-G LV transduction. Additionally, we examined the impact of CAR molecules on the transgene expression and studied the effect of NK cell freezing on transduction efficiency. This study presents a novel combination of BX795 and Retronectin to enhance lentiviral transduction of NK cells without compromising cell viability, phenotype or expansion.

## Results

### NK cells are successfully transduced with VSV-G-pseudotyped LVs carrying GFP

To better understand the surface expression of LDLR on freshly isolated and on activated NK cells, we analysed LDLR expression using flow cytometry (Fig. 1A-B). First, we examined freshly isolated, non-activated NK cells and then analysed them after a three-day activation with IL-2 (500 IU/ml) and IL-15 (140 IU/ml). Our analysis confirmed that freshly isolated NK cells do not express LDLR on their surface, whereas 45% of activated NK cells did. To explore whether rosuvastatin could further enhance the LDLR expression, we treated the IL-2/IL-15-activated cells with rosuvastatin (5 μM). Remarkably, this resulted in expression of LDLR in up to 70% of the cells. The median fluorescence intensity (MFI) increased 10-fold following IL-2/IL-15 treatment, and 20-fold when IL-2/IL-15 was combined with rosuvastatin (Fig. 1A), suggesting an increase in both the number of LDLR-expressing cells and the number of LDLR molecules on each expressing cell.

**Figure 1.**
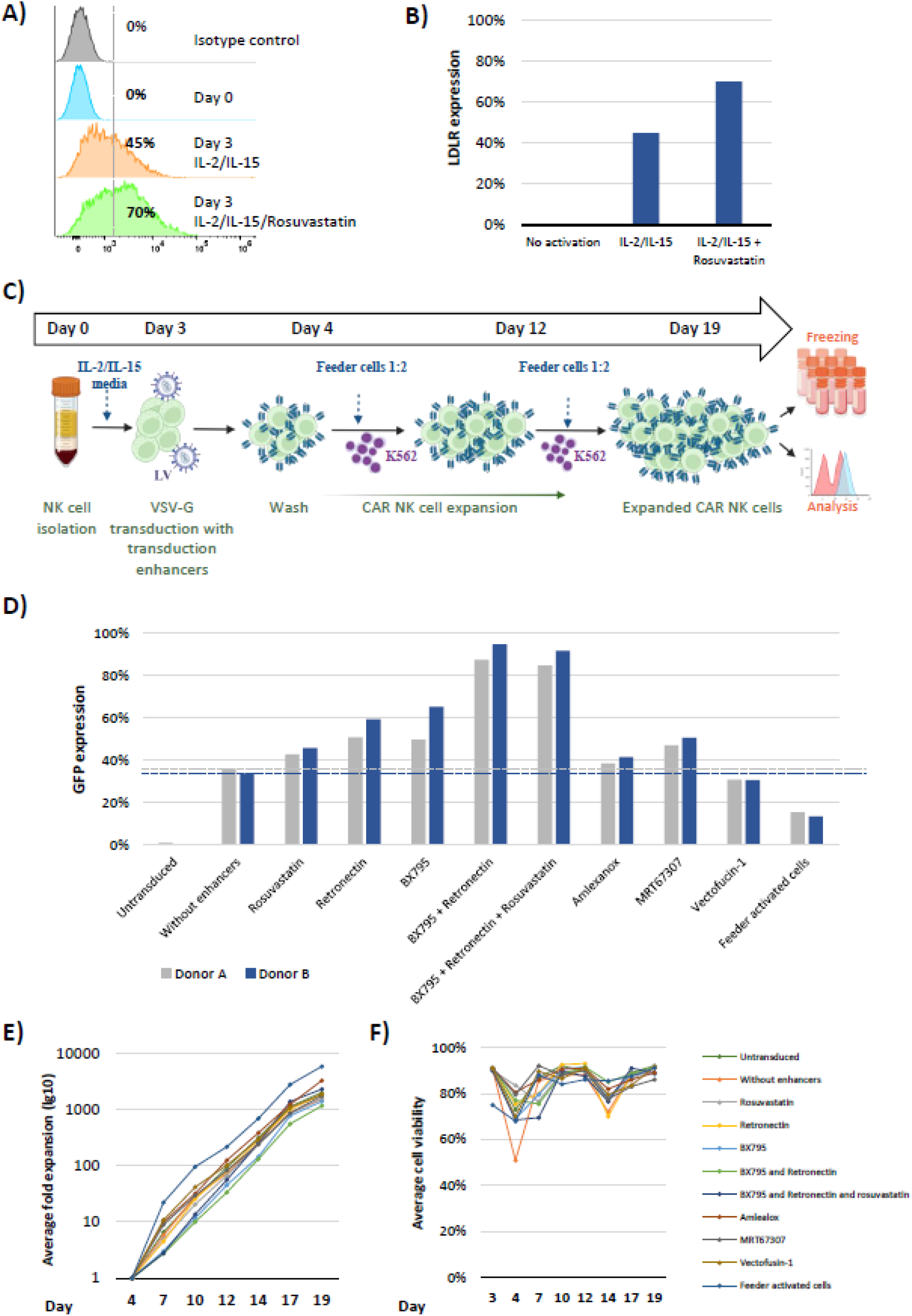
VSV-G GFP transductions of NK cells. A-B) LDLR expression on surface of non-activated, IL-2/IL-15-activated and IL-2/IL-15/rosuvastatin-activated NK cells. The cells were activated for three days and surface expression of LRLD assessed by flow cytometry. N=1 C) The General workflow followed in transducing and expanding NK cells with GFP or CAR carrying VSV-G-pseudotyped LVs. Prior to transduction, NK cells were activated for three days in the presence of IL-2 and IL-15, after which (on day 3) cells were transduced using transduction enhancers. Enhancers and LV were removed by PBS wash on day 4. Total expansion lasted from day 4 until 19 and included addition of irradiated K562 based feeder cells on days 4 and 12. After expansion CAR-NK cells were subjected to analyses or frozen in liquid nitrogen. The figure was created with BioRender.com. D) GFP expression by transduced NK cells. Prior to transduction the NK cells were treated with different transduction enhancers, as listed under the X-axis. GFP expressions of day 19 CAR-NK cells was measured by flow cytometry and shown for individual cultures as columns. Analysis was conducted using two biological replicates (blue and grey). Dashed lines display the level of GFP expression in non-treated NK cells. E) Fold expansion of NK cells after transduction between days 4-19. Fold change (logarithmic) is shown on the Y-axis and fold expansion for each transduction method is depicted with color-coded lines (keys in right panel). Data points show the average expansion of cells from two donors. F) Average viability of NK cells from two donors transduced with GFP using different transduction enhancers. The cells were stained with Trypan Blue and counted with a cell counter. The percentage of viable cells (average of two biological replicates) is shown on the Y-axis and viability curves for individual transduction methods are color coded (keys in right panel).

Next, to compare the efficacy of rosuvastatin, Retronectin, BX795, amlexanox, MRT67307 and Vectofucin-1 in enhancing NK cell transduction, we transduced primary NK cells from two donors with VSV-G-pseudotyped LV carrying the GFP transgene using these enhancers. The transduction workflow includes a two-week, post-transduction, feeder cell-induced CAR-NK cell expansion (Fig. 1C). The efficacy of each agent was studied separately; some of them, having different mechanisms of action, were subsequently combined to interrogate possible synergism (Fig. 1D). For example, inhibition of intracellular TBK1/IKKε antiviral signalling by BX795 was combined with co-localization of target cells with LVs facilitated by Retronectin. This combination was then supplemented with rosuvastatin, which enhances the cell contact further via a complementary mechanism, i.e. upregulating LDLR expression.

Before all transductions, NK cells were pre-activated with IL-2 and IL-15 for three days. Under these conditions, the basal transduction efficiency was 36% and 34% for donors A and B, respectively (Fig. 1D). Of the individual transduction enhancement methods rosuvastatin (donor A: 43% and B: 46%), Retronectin (51% and 59%), BX795 (50% and 65%), amlexanox (38% and 41%), and MRT67307 (47% and 51%, respectively) improved the transduction efficiency, while Vectofucin-1 (31% and 31%) and feeder cell-induced activation (15% and 13%) lowered it. Additionally, a novel combination of BX795 and Retronectin resulted in a synergistic enhancement with a transduction efficiency of (87% for donor A and 95% for B. A triple combination of BX795, Retronectin and rosuvastatin resulted in similarly high levels of transduction (85% for donor A and 92% for B).

In addition to GFP expression, the subsequent expansion (Fig. 1E) and viability (Fig. 1F) of the NK cells were followed during a feeder cell-induced cell expansion. While NK cells activated with feeder cells prior to transduction expanded most efficiently, their low transduction efficiency rendered the method unsuitable for further development. The growth curves of all other cultures receiving feeder cells after transduction were similar to each other, with BX795 slightly diminishing the cell expansion (Fig. 1E).

Cell viability dropped from a mean of 89% (Day 3) to a mean of 72% following transduction (Day 4, Fig. 1F) but recovered by day 10. After the addition of a second batch of feeder cells (on Day 14), the viability decreased in almost all cultures, reflecting the presence of dying feeder cells. However, at the end of the culture period all cultures had good viabilities ranging from 86% to 92% suggesting that the transduction enhancers did not have a major negative effect on NK cell expansion or viability. These results suggest that combining BX795 with Retronectin with or without rosuvastatin yields robust and reproducible transduction of NK cells.

### NK cells are successfully transduced with VSV-G-pseudotyped LVs carrying CD19-targeting CARs

After establishing the most effective, single-agent transduction enhancers and their potent combinations using VSV-G LVs carrying GFP, we investigated whether they could be used to enhance transduction with VSV-G LVs encoding CARs (Fig. 2A). In this analysis, we used the CD19-targeting FiCAR-NK1 construct, the structure of which is illustrated later in Figure 4A. Cells were produced as described above (Fig. 1C) and the CAR expression analysed on day 20. As with VSV-G GFP, BX795 in combination with Retronectin resulted in a high transduction efficiency: 92%, 72% and 77% of NK cells of donors A, B and C expressed a CAR on their surface, respectively. Adding rosuvastatin to this combination did not significantly (p=0.656) improve the transduction efficiency. As before, the transduction enhancers did not markedly impact the feeder-based cell expansion (Fig. S1A). Although cell viabilities varied slightly during the expansion, CAR-NK cells had similar viabilities at the end of the process on day 20 (93-95%, Fig. S1B).

**Figure 2.**
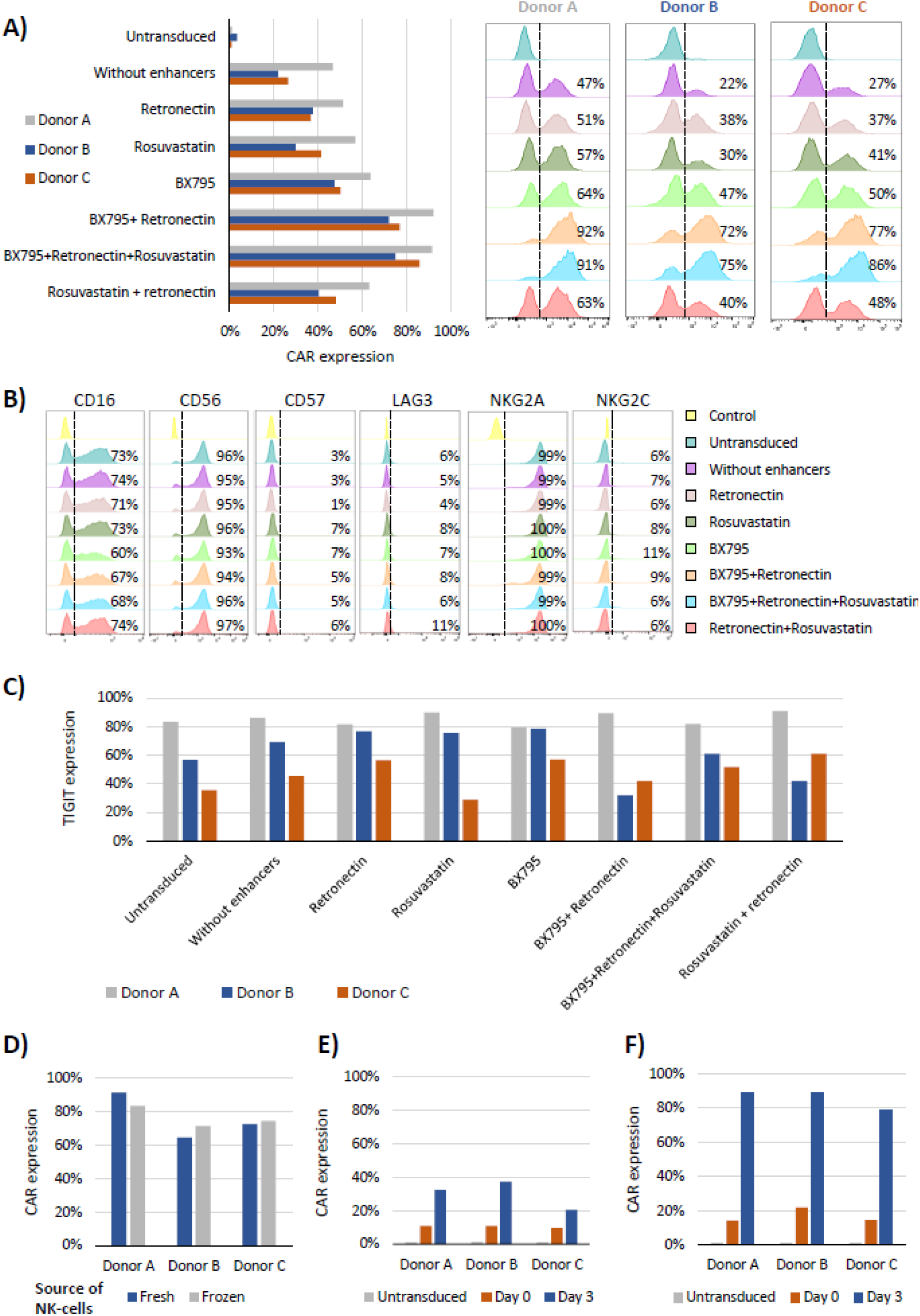
Flow cytometric characterization of CAR-NK cells produced using different transduction enhancers. A) CAR expression of the CAR-NK cells. The different transduction enhancers are listed on the Y-axis. CAR expression of day 19 CAR-NK cells was measured by flow cytometry and shown for individual cultures as columns. Analysis was conducted using three biological replicates (donors). Histograms of the flow cytometry data for each enhancer and three biological donors are shown on the right. B) Representative example of surface expression of selected NK cell receptors (donor A). Data for donors B and C are shown in supplemental figure S1. Receptor expression was analysed by flow cytometry from day 21 CAR-NK cells. Colour coding of different transduction enhancement methods is shown on the right. C) TIGIT expression showed high donor-dependent variation. Percentages of TIGIT positive cells are shown on the Y-axis and different transduction enhancers for each donor (n=3) on X-axis. TIGIT expression was analysed by flow cytometry from day 21 CAR-NK cells. D) Comparison of fresh and frozen NK cells as starting material. Comparison of CAR expression from CAR-NK cells produced from fresh (non-frozen) or frozen-thawed NK cells. Analysis was conducted using three biological replicates (donors). E-F) NK cells were transduced without transduction enhancers (E) or using BX795/Retronectin combination (F) either directly after isolation (day 0) or three-day activation in the presence of IL-2 and IL-15 (day 3). CAR expression was analysed by flow cytometry from day 10 CAR-NK cells. Analysis was conducted using three biological replicates (donors).

To investigate the effect of different transduction methods on cell phenotype, we analysed the surface expression of several phenotypic markers, including activating and inhibitory receptors and exhaustion markers (CD16, CD56, CD57, LAG3, NKG2A, NKG2C and TIGIT) on the CAR-NK cells after 21 days of expansion using flow cytometry (Figs. 2B-C and S1C). Aside from natural variation between donors, the cell phenotypes remained consistent across different transduction methods. For example, TIGIT expression varied markedly between donors but was not affected by the methods employed (Fig. 2C).

Taken together, these data indicate that the combination of BX795 and Retronectin with or without rosuvastatin yields high transduction efficiency without compromising cell expansion, viability, or NK cell phenotype. However, rosuvastatin did not offer a significant added benefit and differences between these two combinations were largely a result of donor-dependent variation (Fig. 2A). To simplify the transduction protocol and make it more suitable for clinical applications, we selected the combination involving only two chemicals, BX795 and Retronectin, for further experiments.

In the previous set of experiments, transduction was performed on freshly isolated and activated NK cells. To determine whether similar results could be obtained using thawed NK cells frozen directly after isolation, we transduced NK cells from three donors either directly after isolation or after freezing and thawing and analysed the CAR expression on day 20 (Fig. 2D). No significant difference (p=0.96) was seen in the transduction efficiency between the fresh or frozen NK cells (Fig. 2D), indicating that both may be used as starting material. However, a fraction of NK cells is lost during freezing and thawing potentially reducing the amount of starting material.

We then examined whether pre-activation of NK cells with IL-2 and IL-15 (Fig. 1C) is necessary for efficient transduction. Freshly isolated NK cells were transduced either immediately after isolation or after a three-day activation with IL-2 and IL-15. Transduction was performed without enhancers (Fig. 2E) or with a combination of BX795 and Retronectin (Fig. 2F). The resulting CAR expression was low (10-11% on day 10) on non-activated NK cells transduced without enhancers (Fig. 2E). Pre-activation increased the CAR expression to 20-37%. Transduction of non-activated NK cells with BX795 and Retronectin resulted in 14-22% CAR expression, whereas pre-activated cells transduced with enhancers had the highest transduction efficiency, resulting in a CAR expression of 79%-89% (Fig. 2F). This suggests that both NK cell pre-activation and the use of transduction enhancers are required to acquire high transduction efficiency.

### Proteomic analysis reveals minimal changes between differentially transduced NK cells

After establishing efficient transduction conditions for NK cells, we employed proteomic analysis by mass spectroscopy to assess potential alterations in the cells resulting from the transduction enhancement. NK cells from three donors were transduced with VSV-G-pseudotyped LV carrying the CD19 targeting CAR construct (FiCAR-NK1). Activated NK cells were transduced either without enhancers or using the combination of BX795 and Retronectin. Untransduced NK cells served as controls. We evaluated the CAR expression using flow cytometry (Fig. S2A) and assessed the functionality of the CAR-NK cells through a cytotoxicity assay against RS4.11 cells (Fig. S2B). For proteomic analysis, we collected 10 x10^6^ unsorted CAR-NK cells (day 19) in four technical replicates. The cells were lysed and digested with trypsin into peptides. Half of the resulting peptide mixture was analysed for total protein content by liquid chromatography mass spectrometry (LC-MS), while the other half underwent enrichment of the phosphorylated peptides using Ti^4+^ beads prior to analysis by LC-MS.

The total protein analysis identified and quantified 5727 most abundant proteins expressed in the (CAR-)NK cells (Table S1A). To gain a more comprehensive understanding of protein expression variation across different sample groups, we conducted a PCA analysis which uncovered greater variation between the donors than among differentially transduced NK cells (Fig. 3A). We did, however, notice slight subclustering within individual biological donors based on different conditions (untransduced, transduced without enhancer, or transduced with BX795 and Retronectin). Yet the order of these subclusters in the PCA plot did not align consistently across different donors. Hierarchical clustering yielded similar outcomes: biological donors formed distinct clusters, and within each donor, we observed subtle subclustering based on different conditions (Fig. 3B). Taken together, these findings indicate that the variation in protein expression between biological donors far outweighs the variation attributed to the different transduction methods.

**Figure 3.**
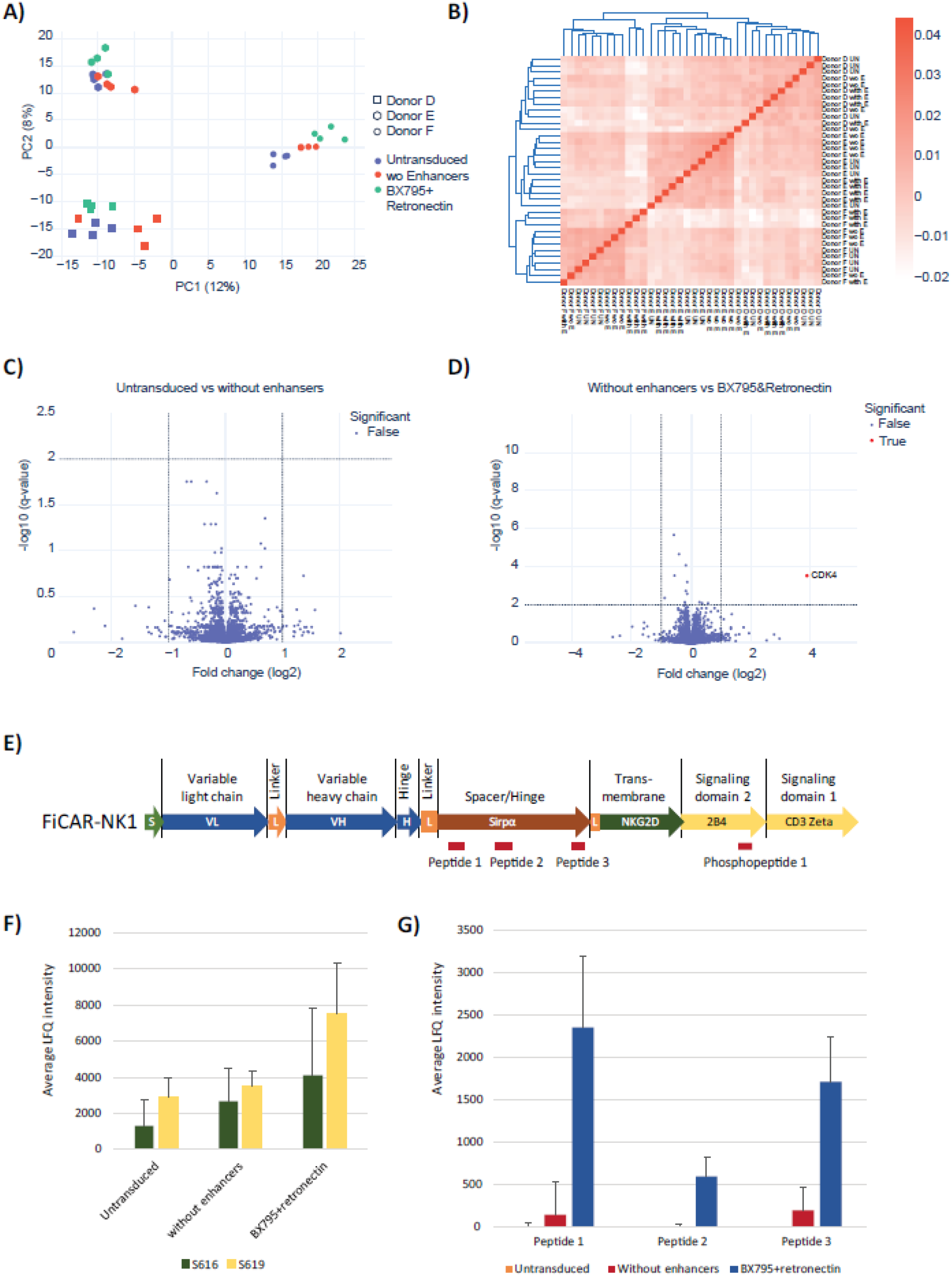
Proteomic analysis of CAR-NK cells produced using BX795 and Retronectin. A) Principal compound analysis (PCA) was performed for the protein expression levels of three different donors and three different treatment groups. Different transduction enhancer conditions are color-coded, and different donors illustrated with different symbols (keys on right panel). Analyses were performed with three biological replicates (donors) each of which had four technical replicates. B) Hierarchical clustering of protein profiles of three different donors and different treatment conditions. C-D) Volcano plots illustrating non-significant (blue dots) and significant (red dots; fold change >2 and −log10 q-value >2) changes in the NK cell proteome. Plot C compares untransduced NK cells to those transduced without enhancers, while plot D compares cells transduced without enhancers to those treated with BX795 and Retronectin. E) Schematic overview of domain organization of the FiCAR-NK1 construct used in the proteomic analyses. The specific locations of the detected peptides and phosphopeptides are indicated below the schematic of the construct. D) Average label free quantifications (LFQs) of CD288 (2B4) phophopeptide in differentially transduced NK cells. Peptide was phosphorylated in serine 616 (S616, dark green) or in serine 619 (S619, yellow) and the corresponding LFQs are shown as own columns. E) LFQs of detected SIRPα/SIRPβ1 peptides 1-3 in differentially transduced NK cells. Proteomic and phosphoproteomic analyses were conducted using three biological replicates (donors), each of which with four technical replicates. Data in F-G are represented as average ± SEM.

To analyse the differences in protein expression on a more detailed level, we calculated the q-values and fold changes for each protein or protein group across different sample groups and visualized them using volcano plots (Fig. 3C-D). A comparison between the untransduced cells and cells transduced without enhancers did not reveal any significantly differentially expressed proteins (fold change >2 and −log10 q-value >2; Fig. 3C, Table S1B). This indicates that lentiviral infection alone has no impact on the expression of these 5727 proteins after a two-week expansion following transduction. Next, to assess the impact of BX795 and Retronectin on the proteome of transduced NK cell, we compared cells transduced without enhancers to samples transduced with BX795 and Retronectin (Fig. 3D, Table S1C). This analysis revealed only one differentially expressed unique protein (fold change >2 and −log10 q-value >2). Out of 5727 proteins, only the expression of CDK4 was significantly lower in BX795 and Retronectin-treated samples compared to samples transduced without enhancers.

While the differentially transduced NK cells appear to express similar levels of the over 5700 most abundant proteins, the resulting protein identifications serve as a valuable resource for understanding the protein expression in the feeder cell-expanded CAR-NK cells (Table S1). To gain deeper insights into the CAR-NK cell proteome, we conducted several enrichment analyses on the proteins identified. These included Biological Process Gene Ontology (GO-BP) (Table S1D), KEGG pathway enrichment (Table S1E), and REACTOME pathway enrichment analysis (Table S1F).

The GO-BP analysis revealed enrichment in several pathways related to mRNA processing, protein transport, cell cycle, and translation (Table S1D). KEGG analysis highlighted enrichment of various pathways, including metabolic pathways (p=2.8E-15), oxidative phosphorylation (p=4.98E-11), Fc gamma R-mediated phagocytosis (p=5.7E-05), and natural killer cell-mediated cytotoxicity (p=2.5E-04, Table S1F). A closer examination revealed that most proteins of the natural killer cell-mediated cytotoxicity pathway were detected in our analysis (Fig. S2C). These proteins include, for example, all the NK cell surface receptors relevant to the pathway.

Given that BX795 is known to impact the intracellular signalling pathways of NK cells, we conducted a Ti4+ bead-based phosphoprotein enrichment followed by LC-MS analysis of the phosphorylated proteins. This analysis identified a total of 1957 phosphorylated proteins (Table S2A). Notably, we observed no significant difference in the expression of these phosphorylated proteins between the untransduced and enhancer-free transduced NK cells (Table S2B) suggesting that the viral infection itself did not affect the NK cell phosphoproteome. However, two proteins, ENSA and CD244 (also known as Natural Killer cell receptor 2B4), showed elevated phosphorylation in BX795/Retronectin-treated compared to enhancer-free transduced samples (Table S2C). Interestingly, the CAR construct utilized in our experiments, (CAR-NK1, Fig. 3E), incorporates a signalling domain originating from 2B4. This led us to investigate whether higher expression of 2B4 could be attributed to an increased CAR expression on NK cells. A detailed examination of the enriched phosphorylated 2B4 revealed the presence of a singular phosphopeptide whose sequence aligns with the 2B4 signalling domain incorporated within the CAR construct (Fig. 3E). Notably, this phosphopeptide contains two phosphorylation sites, positions S616 and S619. Both sites exhibited augmented phosphorylation levels in cells treated with BX795 and Retronectin (Fig. 3F).

Finally, a total protein analysis identified three peptides corresponding to the signal-regulatory protein alpha (SIRPα)-derived spacer segment of the CAR construct (Fig. 3E). The expression levels of these peptides were also elevated in the BX795/Retronectin -treated cells (Fig. 3G). Of note, these peptides are not exclusive to SIRPα but also shared with SIRPβ1 and were consequently excluded from the total protein analysis. Nevertheless, these findings imply that an increase in CAR expression results in the detection of the CAR domains in the proteomic analysis.

### Combination of Retronectin and BX795 enhances transduction of different CD19-targeting CARs

Next, we asked whether CARs with varying structures can be expressed on NK cells with equal efficiency. Earlier, we have used the FiCAR construct to successfully modify T cells.^16^ The FiCAR backbone contains an extracellular spacer derived from the immunoglobulin-like domains of SIRPα and allows for the design of multiple structural variants of the CAR.^17^ For this study, we designed three novel NK-specific, CD19 targeting CAR constructs carrying signalling domains from 2B4 and CD3z involved in activating signal transduction of NK cells. Two of these novel constructs, FiCAR-NK1 and FiCAR-NK2, contain the SIRPα spacer (Fig 4A). The third, CAR-NK3, contains a CD8 alpha-based spacer (Fig. 4A).

**Figure 4.**
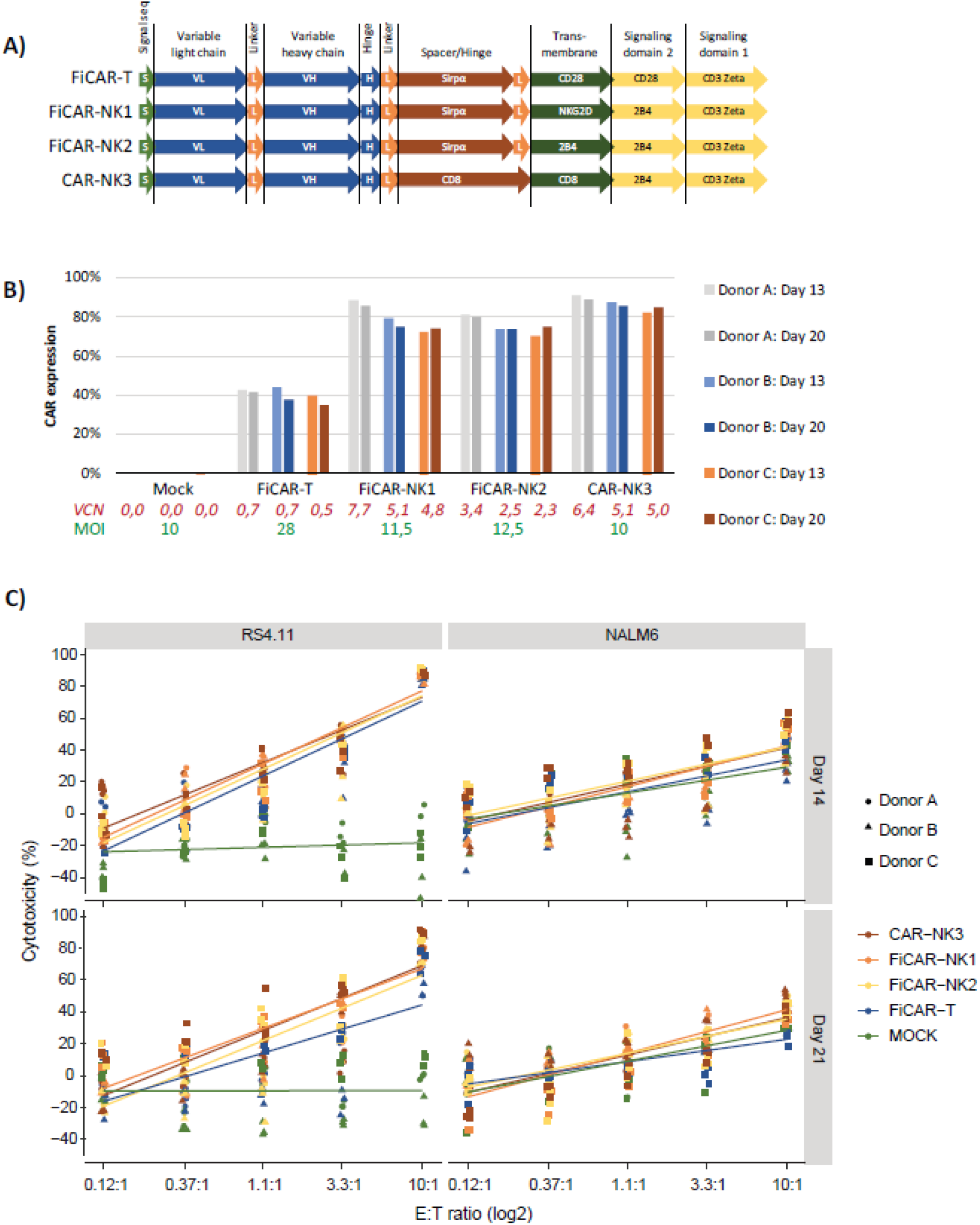
Transduction of CARs with different structures. A) Schematic illustration of the domain organisation of the CAR-constructs used in this analysis. B) Percentage of CAR-expressing NK cells derived from three different biological donors (n=3) on day 13 (light colors) and on day 20 (dark colors). Vector copy numbers (VCN) from day 19 CAR-NK cells and the multiplicity of infection (MOI) ratios are shown below the X-axis using red and green fonts, respectively. C) Cytotoxicity of different CAR-NK cells against RS4.11 and NALM6 ALL cell lines. CAR-NK cells were incubated with target cells for four hours. Linear regression was performed on the log2 transformed E:T ratios per construct group (three biological donors with three technical replicates). Donors are highlighted with different shapes (key in right panel) and CAR constructs are color-coded. Cytotoxicity was assayed on day 14 (upper graphs) and 21 (lower graphs). The effector to target ratios (E:T) are shown on the X-axis.

As before, NK cells were pre-activated with IL-2 and IL-15, and the BX795/Retronectin combination was used to enhance transduction. To achieve comparable CAR expression for functional assays, varying amounts of vectors were used for each construct. The multiplicity of infections (MOIs) for each construct was selected based on CAR expression from previous experiments (data not shown) and is illustrated in Figure 4B. All vectors were produced, and the virus titers were measured at the same time (Table S3). Transduction with the combination of BX795 and Retronectin resulted in a high transduction efficiency and expression of NK-specific CAR-constructs on the cell surface (Fig. 4B). Interestingly, the expression of T cell specific FiCAR-T on NK cells was significantly lower (p<0,001) compared to the expression of NK-specific CAR constructs despite an MOI more than two-fold higher (Fig. 4B). This was unexpected as the same FiCAR-T LVs have yielded high transduction efficiency in T cells.^16^

To investigate whether CAR expression changes during cell expansion, the surface expression was analysed by flow cytometry on days 13 and 20 of the cell production. The percentage of CAR expressing cells remained stable during the NK cell expansion (Fig. 4B) indicating that the expression neither favoured nor inhibited the expansion of transduced cells over non-transduced cells.

Next, we studied the cytotoxicity of the different CAR-NK cells using a luciferase-based cytotoxicity assay. CAR-NK cells (on days 14 and 21) were cocultured with luciferase expressing acute lymphoblastic leukemia (ALL) target cells (RS4.11 or NALM-6) for four hours and the remaining live target cells quantified by luminescence (Fig. 4C). The RS4.11 target cells were highly resistant to the natural cytotoxicity of unmodified NK cells and only killed by the CAR-NK cells, whereas NALM-6 cells were killed equally well by MOCK-and CAR-NK cells. NK cells transduced with NK-specific CARs induced similar cytotoxicity, indicating that the FiCAR-based, NK-specific CARs (FiCAR-NK1 and FiCAR-NK2) are equally cytotoxic compared to the CD8-based CAR-NK3. On day 21, FiCAR-T transduced NK cells, however, were slightly less cytotoxic against RS4.11 target cells than cells transduced with an NK-specific CAR. On day 14 and 21 FiCAR-T-NK and MOCK-NK cells showed impaired cytotoxicity against NALM-6 at high effector:target ratios (3,3:1 and 10:1).

To demonstrate the expected donor-dependent variation in CAR-NK cytotoxicity, we plotted each donor separately (Fig. S3). Indeed, the target cells were killed more efficiently by different donors. For example, day 21 CAR-NK cells from donor B were more cytotoxic against NALM-6 cells than day 21 CAR-NK cells from donors A and C. Against RS4.11 cells, however, CAR-NK cells from donor B were less cytotoxic than cells from donor A or C. The donor variation was also affected by the age of the cells: donor B showed lowest cytotoxicity against NALM6 cells on day 14 but the highest on day 21.

Taken together, we were able to produce CAR-NK cells with a high transduction efficiency and consistent cytotoxicity using different NK-specific CARs. The expression of the T cell-specific FiCAR-T construct, however, was significantly lower, resulting in slightly impaired cytotoxicity.

### The CAR structure influences VSV-G-LV-mediated transduction in NK cells

To examine whether the lower expression of FiCAR-T compared to that of the NK-specific CARs was the result of lesser integration of the CAR-construct into the genome or impaired expression of the CAR protein from the integrated gene, we analysed the vector copy number (VCN) in the cell product by ddPCR (Fig. 4B). The results show that the VCN obtained with FiCAR-T was much lower (0,5-0.7 copies per cell) than with the NK-specific CARs (2,3-7,7 copies per cell) suggesting a more efficient integration of the NK-specific CAR. Of note, the VCN number is the average across the whole cell batch, and not of the CAR expressing cells only, which prevents direct comparison of VCN between CAR-transduced cells alone.

To better understand the lower transduction efficiency of the FiCAR-T VSV-G LVs, we titrated a high quality, reactor scale version of the FiCAR-T (FiCAR-T(R)) VSV-G LV from a different manufacturer using NK and T cells. FiCAR-T(R) shared the main domain organisation of FiCAR-T but had a small modification in the hinge domain to increase stability (Koski et al. 2025, submitted). First, the VSV-G LV encoding CAR-NK3 was titrated on NK and T cells as a positive control for transduction efficiency. This resulted in comparable levels of CAR expression and vector copy numbers in both NK and T cells (Fig. 5A-B) indicating that the transduction protocols (activation and BX795/Retronectin for NK and activation only for T cells) were functional and efficient at MOI one and higher. Next, using the same transduction protocols and the same donor, the FiCAR-T(R) LV was titrated using T- and NK cells (Fig. 5A and C). The LV efficiently transduced T cells (73%, MOI 10), whereas NK cell transduction efficiency with Retronectin and BX795 remained low (43%, MOI 500). Additionally, the vector copy numbers were approximately three times higher in T compared to NK cells. Similar results were obtained when FiCAR-T(R) was titrated using the NK and T cells from different donors (Fig. S4). These results suggest that the amount of expanded NK cells expressing a functional CAR is dependent on both the transduction method and the CAR expressed.

**Figure 5.**
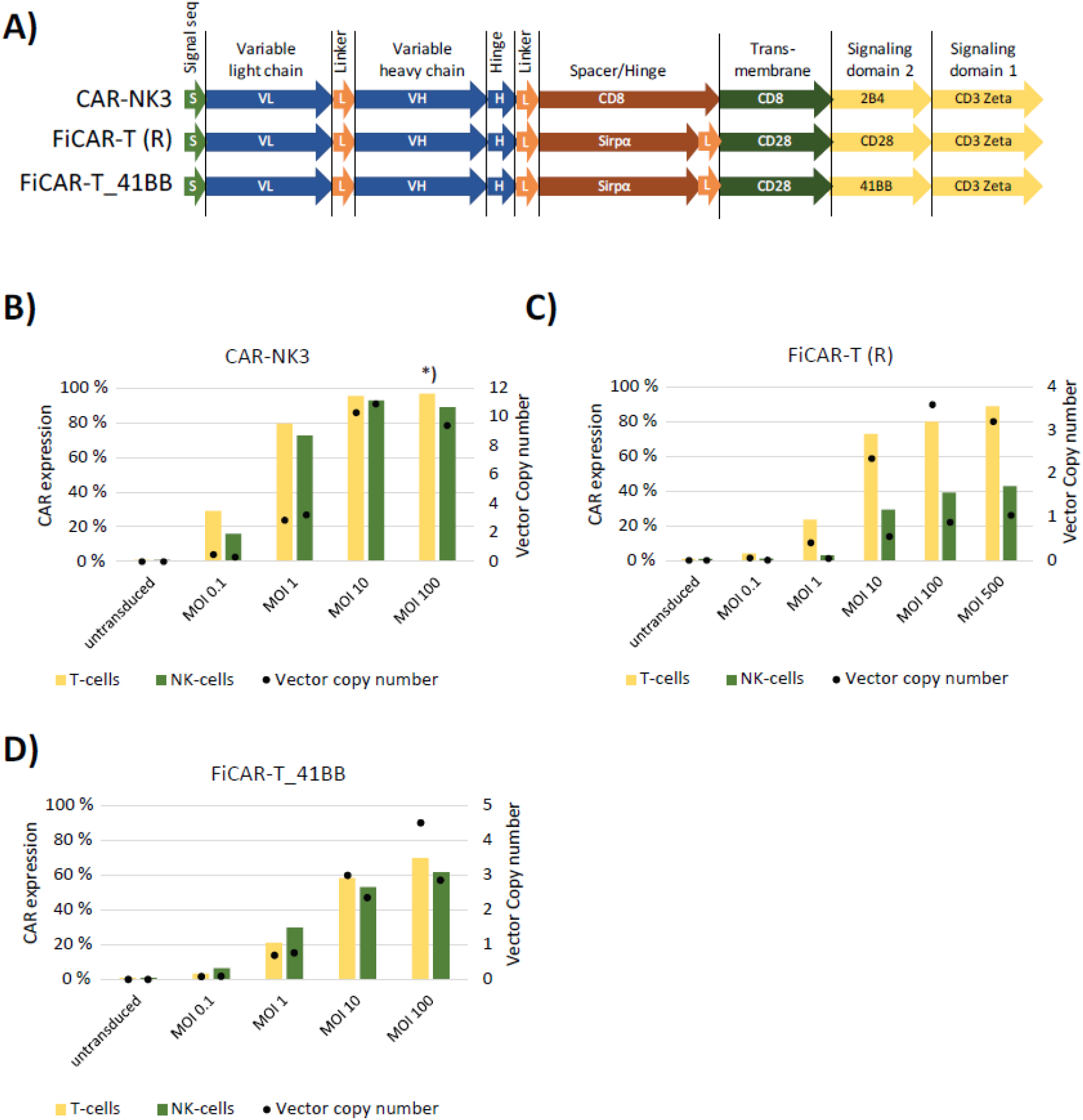
The effect of the CD28 and CD4-1BB signalling domains on transduction efficiency. A) Schematic illustration of the domain organisation of the CAR-constructs. FiCAR-T (R) was similar to FiCAR-T used previously (Figure 4), albeit having a minor modification in the hinge domain. In FiCAR-T_41BB the CD28 signalling domain 2 was replaced by one from 4-1BB. CAR-NK3 was used as positive control for transduction conditions. B-D) CAR expression and vector copy number in NK and T cells after transduction with titrated LV vector encoding (B) CAR-NK3, (C) FiCAR-T (R) and (D) FiCAR-T_41BB. NK cells were transduced following IL-2/IL-15 activation and using BX795 and Retronectin for enhancement. T cells were activated with Transact-beads prior to transduction. CAR expression was assessed on day 12 for NK cells and on day 9 for T cells by flow cytometry.

The only structural differences between the FiCAR-T and FiCAR-NK1/NK2 constructs are the transmembrane and signalling domains, which in the FiCAR-T are derived from CD28 and in NK-CARs from NKG2D, 4B2 and CD8 (Fig. 4A). This suggest that the differences in CAR expression are caused by the CD28 domains. To further study whether the CD28 signalling domain causes a reduced transgene expression, we transduced NK- and T-cells with a FiCAR-T construct in which the signalling domain was replaced with one from 41BB (FiCAR-T_41BB; Fig. 5A). The substitution led to similar CAR expression on both NK and T cells with slightly higher vector copy numbers in T cells (Fig. 5D). This further supports the conclusion that the impaired transduction and expression of T cell-specific CARs on NK cells is indeed caused by the CD28-derived signalling domain.

### Baboon envelope-pseudotyped LV transduction shows fluctuation in virus titers and CAR expression

Recently, Baboon envelope-pseudotyped lentiviral vectors (BaEV LVs) have been successfully used for NK cell transduction.^11,18^ Upon entering its target cell, the BaEV LV binds to the ASCT2 receptor expressed on activated NK cells.^11^ To determine the optimal day for transduction, we screened the expression of ASCT2 upon IL-2/IL-15 activation by Western blot (Fig. 6A). Based on the ASCT2 expression, day 3 was selected for transduction. For the initial experiments, we produced a BaEV LV carrying GFP and transduced NK cells using two enhancement methods, Vectofusin-1 and spinoculation (Fig. 6B). Both Vectofusin-1 and spinoculation enhanced transduction, and by using both, we could achieve a nearly 100% transduction efficiency with a MOI of 5. However, compared to the VSV-G LV production, the titers of the BaEV LVs were on average 2 logs lower (Table S3, Fig. S5).

**Figure 6.**
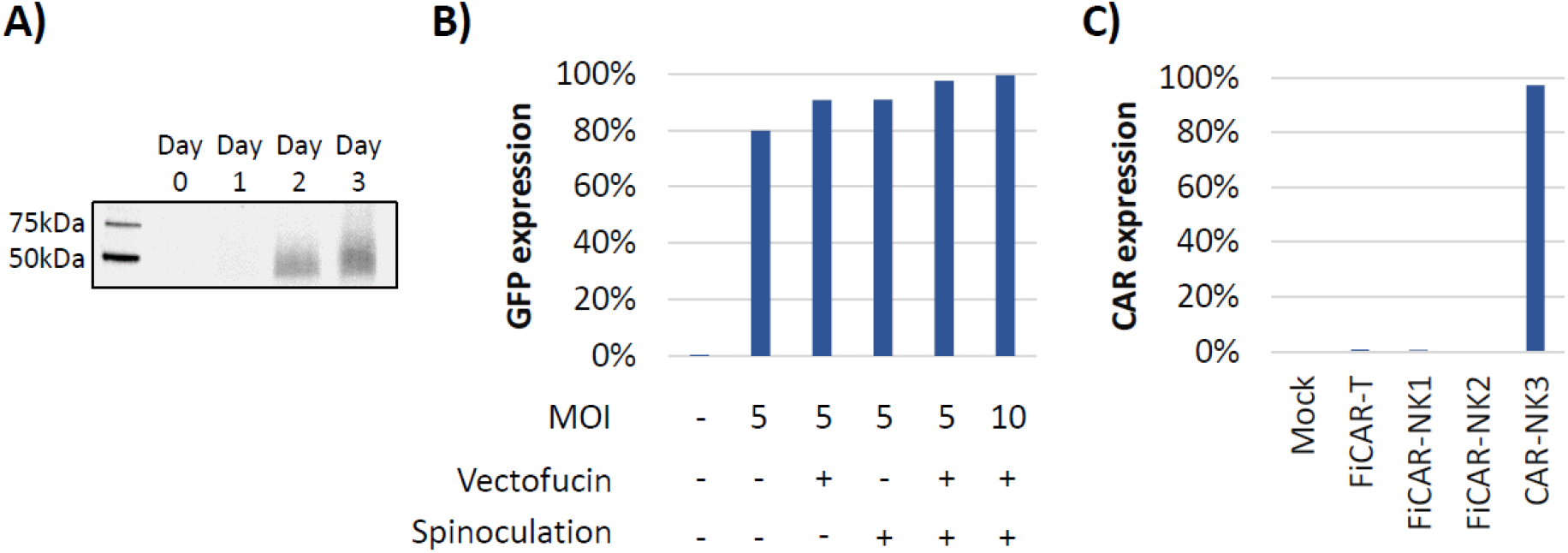
BaEV -LVs in NK cell transduction. A) ASCT2 expression in NK cells during activation with IL-2/IL-15. Gel staining for loading control before anti-ASCT2 labeling is shown in supplemental figure S6. B) GFP expression (%) after transduction eith BaEV pseudotyped virus encoding GFP. The MOI ratio, and use of Vectofusin-1 and spinoculation are shown under the X-axis. C) CAR expression after transduction with BaEV pseudotyped virus encoding different CARs. BaEV-LVs were produced using same transfer plasmids as those used in VSV-G-pseudotyped LV production.

To test the usefulness of BaEV LVs for the transduction of FiCAR-T, FiCAR-NK1, FiCAR-NK2 and CAR-NK3 constructs, we produced the lentivectors using the same transfer plasmids as for the VSV-G production, only substituting the BaEV envelope plasmid for the VSV-G plasmid. Surprisingly, only the CAR-NK3 carrying BaEV lentivectors were functional resulting in a high transduction efficiency (97%, MOI 5) (Fig. 6C). This was unexpected as the same transfer plasmids were functional in the VSV-G production and the production process of the LVs was successful as both the GFP and CAR-NK3 BaEV LVs were functional. The LV production process was repeated with identical results: functional GFP and CAR-NK3 BaEV LVs were produced, but the BaeV LVs carrying FiCAR-T and FiCAR-NK1/NK2 genes failed to transduce NK cells under identical conditions. This may indicate that the CAR construct composition plays a role in BaEV LV production and/or transduction. Subsequently, due to low titers and mixed transduction results, experimentation using BaEV-pseudotyped LVs was discontinued for the present study.

### Retronectin and BX795 do not enhance T cell transduction

Finally, given the potency of Retronectin and BX795 to enhance the VSV-G lentiviral transduction of NK cells, we were curious to learn whether the combination is also effective for the transduction of T cells. While the genetic modification of T cells can readily be accomplished using current methodology, any improvements in efficiency could yield cost savings in therapeutic cell production. To answer this question, T cells were transduced with VSV-G FiCAR-T(R) without enhancers or with BX795, Retronectin or the BX795/Retronectin combination (Fig. 7A). This analysis showed clearly that BX795 alone or in combination with Retronectin did not enhance T cell transduction but was in fact inhibitory. Retronectin alone had little effect, and the detrimental effect of BX795 may be related to its toxicity on the T cells, as cell expansion was also impaired in cells treated with BX795 (Fig. 7B).

**Figure 7.**
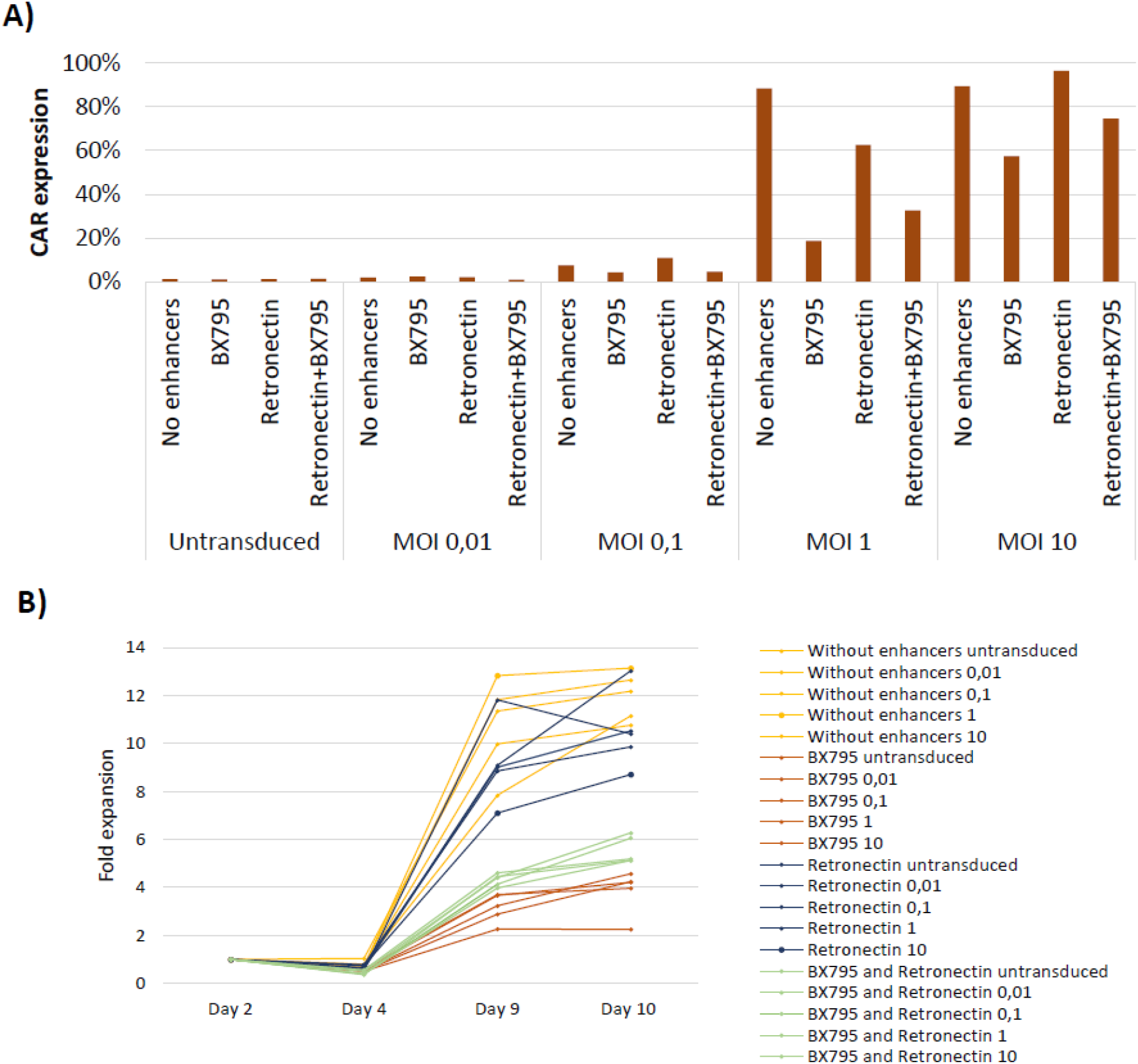
BX795 and Retronectin in T cell transduction. A) T cell specific FiCAR-T (R) was titrated to T cells without enhancers or in the presence of BX795 and/or Retronectin. B) CAR-T cell expansion was monitored after transduction. Cells transduced in the presence of BX795 showed lower expansion compared to those transduced without enhancers or with Retronectin only. MOIs are indicated in the figure legend. N=1.

## Discussion

Recently, genetically modified lymphocytes such as CAR-T and -NK cells have garnered growing interest in cancer cell therapy. While some CAR-T cell therapies have already gained marketing approval, CAR-NK cell therapies are still in preclinical and clinical development. γ-retroviruses and VSV-G-pseudotyped lentiviruses have been the traditional ‘workhorses’ for genetic modification of T cells. Although both approaches have a good safety record in the manufacture of therapeutic immune cells, the former may carry more risk, perhaps due to a different integration profile into the target cell genome.^19^ While efficient transduction of NK cells has been achieved using γ-retroviruses,^20^ transduction with VSV-G-pseudotyped LVs is challenging.^11^ To address this gap, we have optimized the use of the safe and widely used VSV-G-pseudotyped LVs for NK cell transduction.

NK cell transduction with VSV-G LVs can be enhanced through three main mechanisms. First, by NK cell activation and increasing the expression of the natural virus receptor, LDLR, prior to transduction. Second, by facilitating the contact between the LV and NK cells either by promoting the colocalization of viral particles and NK cells or by neutralising the electrostatic repulsion between them. Third, by inhibiting the intracellular antiviral signalling induced by VSV-G infection in NK cells. Our study explored various approaches of activating NK cells and increasing the expression of the LDLR (with IL-15, IL-2, rosuvastatin and feeder cells), facilitating the virus-cell contact (Retronectin, Vectofusin-1) and influencing antiviral signalling (including Amlexanox, MRT67307 and BX-795).^10,12–15^ We investigated these approaches as either single agents and in different combinations aiming to optimize transduction efficiency of VSV-G LVs in NK cells.

Gong et al. demonstrated that Vectofusin-1 did not significantly enhance transduction of primary NK cells by VSV-G LVs which concurs with the findings in this study.^10^ Our experiments revealed that Retronectin outperforms Vectofusin-1 in transduction enhancement (Fig. 1D). However, a recent comparison of Retronectin and Vectofusin-1 for transduction of primary NK cells with a VSV-G-pseudotyped LVs indicated that they have similar efficacies.^21^ In addition, Vectofucin-1 is used in large scale CAR-NK cell production using the CliniMACS Prodigy® platform for clinical studies.^22^ In our experiments, Vectofucin-1 enhanced the transduction of primary NK cells with BaEV-pseudotyped LV (Fig. 6B). Interestingly, when the NK92 cell line was used, Vectofucin-1 completely blocked transduction (data not shown). These results indicate that the effectiveness of Vectofucin-1 depends on the LV-pseudotype used, the specific cell line employed and possibly the method of transduction.

Rosuvastatin has been reported to significantly enhance the transduction efficiency of NK cells.^10^ In our experiments, using NK cells activated with IL-15 and IL-2, we observed a similar improvement, as shown in Figures 1D and 2A. The mechanism underlying the improved efficiency involves upregulation of the LDLR, as demonstrated by our findings (Fig. 1A-B) and others.^10^

NK cell transduction can also be enhanced by inhibiting intracellular antiviral signalling, as demonstrated by Chockley et al.^13^ In their study, they targeted the TLR4 signalling pathway: TBK1 using Amlexanox, IKK⍰ using MRT67307, or both TBK1 and IKK⍰ using BX795. The resulting transduction efficiencies were as follows: 36.1% (Amlexanox), 46.5% (BX795), and 47.3% (MRT67307) in IL-2-activated NK cells.

Consistent with the findings of Chockley et al., Amlexanox yielded a lower transduction efficiency (39.5%) in our study. However, contrary to their results, BX795 led to the highest transduction rates (57.5% using GFP and 53.8% using CAR) and was therefore selected for further experiments (Fig. 1D). BX795 has also previously been reported to enhance NK cell transduction, with efficiencies varying from 20 to 50%.^23^

Next, we investigated whether combining the most efficacious enhancement methods facilitating virus-cell contact and interfering with intracellular signalling would synergize in enhancing transduction efficiency. Notably, we found that the highest level of transduction was achieved when the TLR4 signalling was impacted through TBK1/IKKε inhibition with BX795, and the virus-cell contact physically facilitated using Retronectin-coated culture plates (Figs. 1D and 2A). The addition of a third enhancer, rosuvastatin, to the combination increased the transduction efficiency slightly, but not significantly. By combining BX795 and Retronectin we could achieve a high transduction efficiency: 91% using GFP and 80% using CAR, (averages of two and three biological replicates, respectively). This BX795 and Retronectin combination outperformed transduction efficiency reported in previous studies, demonstrating its potential for CAR-NK cell therapies.

To investigate the potential effects of transduction enhancers on NK cells, we closely examined the growth, viability, and surface marker expression of the CAR-NK cells following transduction with various enhancers. Notably, cell growth and viability were not markedly affected by the enhancers. Additionally, the expression of surface markers, such as CD16, CD56, CD57, LAG3, NKG2A, NKG2C, and TIGIT, exhibited greater variation among biological donors than between the different transduction methods.

These surface marker expression findings align with the results of the total protein analysis, in which we identified and quantified approximately 5700 highly expressed proteins in differentially transduced CAR-NK cells using an MS-approach (Fig. 3). Specifically, we compared the protein expression profiles of untransduced NK cells to those of cells transduced with or without enhancers (BX795/Retronectin). Interestingly, among the proteins identified, most components of the natural killer cell-mediated cytotoxicity KEGG pathway (Fig. S2C) were identified but showed no significant differences between transduction conditions. In fact, similar to the surface receptor analysis by flow cytometry, the expression of these proteins exhibited greater variability among the donors than between the different transduction methods (Figs. 3B and S1C).

We did not detect significant differences in protein or phosphoprotein expression between the untransduced NK cells and those transduced without enhancers. This suggests that lentiviral transduction itself does not impact the expression of these proteins, many of which are involved in intracellular signalling, after a two-week expansion post-transduction. To investigate whether transduction using the BX795/Retronectin combination affects the protein expression of expanded NK cells, we compared the proteomic profiles of enhancer-free, transduced CAR-NK cells to those transduced using BX795/Retronectin. This analysis revealed that CDK4 exhibited a lower expression in the BX795/Retronectin -transduced CAR-NK cells (Fig. 3D) and, in addition, ENSA and CD244 showed increased phosphorylation (Table S2C). ENSA is a protein phosphatase inhibitor related to cell cycle regulation^24^ (Uniprot.org), while CD244 (also known as natural killer cell receptor 2B4) serves as an activating receptor for NK cells. These results may indicate changes in cell cycle regulation, a finding that should be studied further.

Taken together, as only minimal changes were detected in the NK cell surface markers, proteome and phosphoproteome, we postulate that using BX795/Retronectin in transduction does not significantly impact the final CAR-NK product. Additionally, the resulting protein list from the total protein analysis, along with its various annotations (such as GO-BP, KEGG, and REACTOME), provides a valuable resource for understanding the protein expression patterns in feeder cell expanded CAR-NK cells (Table S1).

Activation with cytokines such as IL-2 and IL-15 is essential for an efficient transduction of NK cells with VSV-G LVs.^23^ However, to improve the cost-effectiveness of therapeutic cell production, transduction should be performed as early as possible to minimize LV usage. Previous studies have shown that a two-to three-day activation period increases transduction efficiency.^23^ Consequently, we chose to transduce the cells on day three. As depicted in Figure 2E-F, activation plays a crucial role in NK cell transduction, and the effectiveness of the BX795/Retronectin combination relies on NK cell activation. Additionally, the results shown in Figure 1A-B suggest that the impact of NK cell activation is likely due to increased LDLR expression in the NK cells.

Cord blood units can be utilized as a source of gene-modified NK cells and are often stored in frozen form.^20^ Both in research and in clinical therapy, the use of frozen NK cells as starting material can be advantageous. To assess whether freezing the NK cells prior to transduction affects transduction efficiency, we isolated cells from three distinct buffy coats and performed transduction on either freshly prepared or freeze-thawed samples. Remarkably, the transduction levels remained consistent, suggesting that frozen NK cells can indeed serve as suitable starting material. However, it’s important to note that a fraction of NK cells is likely lost during the freezing process.

Besides the impact on NK cell activation and the use of transduction enhancers, we discovered that the CAR composition also plays a role efficient transduction or transgene expression by the cells. Surprisingly, CARs successfully used for T cell transduction (FiCAR-T and FiCAR-T(R)) resulted in low integration and expression on NK cells. In contrast, NK cell-specific CAR-NK3 was expressed at high levels both on NK and T cells (Figs. 4B and 5B). Our results suggests that CD28’s signalling domain reduced integration and subsequent transgene expression efficiency in NK but not in T cells, as replacing CD28’s signalling domain with one from 41BB resulted in similar expression in both NK and T cells (Fig. 5D). Unfortunately, our approach did not allow us to dissect whether the inefficiency of transduction was due to the presence of CD28 per se, or to the presence of CD28 within a SIRPa-based CAR structure. It is, however, likely that the mechanism is not due to an inefficiency in the production of SIRPa/CD28-containing LV’s, given that these same virus preparations performed well in the transduction of T cells (Figs. 5 and S4). Furthermore, similar T cell specific SIRPa/CD28-containing CAR-constructs have been used in our previous studied with high transduction efficiency.^16^

When titrated on T cell-specific FiCAR-T(R), the maximum proportion of NK cells expressing a CAR plateaued at approximately 40% (Fig. 5C and Fig. S4). This saturation point was not exceeded, even when the virus:target cell ratio was amplified to a MOI of 500 (Fig. 5C). This observation could suggest that only a certain fraction of NK cells is capable of being transduced with FiCAR-T(R). One explanation might be that the SIRPa/CD28-containing transgene, which is also manifested on the LV membrane, interacts unfavourably with certain NK cell subpopulations. This adverse interaction does not occur with T cells, and could result in inadequate virus entry, integration, and/or CAR expression.

This intriguing phenomenon necessitates further investigation for a more comprehensive understanding. However, we propose that the composition of the CAR influences the efficiency of NK cell transduction. This may partially account for the challenges commonly encountered in NK cell transduction. For a more detailed understanding, it would be beneficial to separately study the CAR positive and negative NK cell fractions. However, this was not the primary focus of this study and was not pursued further.

In addition to VSV-G-pseudotyped LVs, we tested the Baboon envelope -pseudotyped LVs (BaEV LVs) for NK cell transduction.^11,18^ Our results showed that transduction of GFP and CAR-NK3 could reliably be achieved using BaEV LVs. However, the FiCAR constructs containing the SIRPα spacer (FiCAR-T, FiCAR-NK1, and FiCAR-NK2) were not successfully transduced using BaEV, even though the same transfer plasmids were used as in production of VSV-G-pseudotyped LVs. It’s possible that NK cells are more sensitive to different genetic constructs introduced via gene transfer than T cells, but to confirm this, further studies are needed. Due to the low virus titers obtained in BaEV production, we decided not to pursue this approach any further.

In conclusion, this study highlights the challenges and complexities of NK cell transduction, emphasizing the need for careful consideration of various factors, such as NK cell activation, construct design, LV-pseudotype selection, and the use of transduction enhancers to modulate the virus-cell interactions and NK cell anti-viral signalling. By manipulating these elements, effective transduction can be achieved, enabling the use of VSV-G-pseudotyped LVs for therapeutic NK cell production. In our optimized workflow, NK cells are activated with IL-2 and IL-15 followed by transduction with a NK cell specific CAR-construct using VSV-G-pseudotyped LVs in conjunction with BX795 and Retronectin. This approach yields an excellent transduction efficiency without compromising the NK cell phenotype or growth.

## Material and Methods

### Cell lines

RS4.11 (CD19+ acute lymphoblastic leukemia, ALL cell line, CRL-1873™) and NALM-6 (ALL, CRL-3273™) were purchased from ATCC (Manassas, VA, USA). To generate the luciferase-expressing cell lines, RS4.11 cells were transduced with LUC2-EGFP (VB210329-1384uya, (VectorBuilder, Germany) and NALM-6 with the IVISbrite Red F-luc-GFP (RediFect™, Perkin Elmer, Shelton, CT, USA) LVs with MOIs 15 and 10, respectively. After transduction and expansion, the eGFP and GFP positive cells were selected using a Sony SH800 (Sony Biotechnology, San Jose, USA) cell sorter. Both RS4.11 and NALM6 target cells were grown RPMI-1640 medium (Gibco, Thermo Fisher Scientific, Walthan, MA, USA) supplemented with 10% foetal bovine serum (Gibco, Thermo Fisher Scientific, Walthan, MA, USA) and 1% penicillin-streptomycin (Gibco, Thermo Fisher Scientific, Walthan, MA, USA).

To efficiently expand NK and CAR-NK cells, K562 based feeder cells were added into the culture twice at a ratio 2:1. The K562-mbIL-15-41BBL feeder cell line used was generous gift from prof. Dario Campana.^25^ The transduction and expansion protocol are depicted in Fig. 1C.

### CAR structures and lentiviral vectors

The schematic structures of the CAR constructs used in the study are shown in Figures 4 and 5. The CAR designed for making CAR T cells (FiCAR-T) has been previously described (named FiCAR 1; Koski et al 2022). Briefly, all the CAR constructs target CD19 with an scFv derived from the mouse anti-human CD19-targeting antibody FMC63, consisting of the light and heavy chain variable domains connected with a (GGGGS)_4_-linker. Following the ScFv, there is an extended human IgG1-derived hinge and another G_4_S-linker. The extracellular spacer was derived either from SIRPα C1-type 1 & 2 domains (FiCAR-T, FiCAR-T_41BB, FiCAR-NK1 and FiCAR-NK2) or from CD8 (CAR-NK3). The transmembrane domains (TMs) varied between the constructs: FiCAR-T and FiCAR-T_41BB contain a CD28 TM domain, while NK-specific CARs comprise NKG2D (FiCAR-NK1), 2B4 (FiCAR-NK2) or CD8 (CAR-NK3) TM domains. The intracellular signalling domain is derived from CD3 zeta in all constructs and the co-signalling domain from CD28, 41BB or from 2B4.

The transgenes were synthesized and cloned into the lentiviral vector transfer plasmids (Schenkwein et al., 2010) by Genewiz (Azenta Life Sciences, Burlington, MA, USA). The VSV-G- and BaEV-pseudotyped third generation LVs for gene transfer were produced in the National Virus Vector Laboratory at the A.I. Virtanen Institute for Molecular Sciences (University of Eastern Finland, Finland) with standard methods ( https://pubmed.ncbi.nlm.nih.gov/11883085/)^26^ The vectors were characterized for their p24 capsid protein using the Alliance HIV-1 p24 Antigen ELISA Kit (NEK050B001KT, Revvity, Germany). LVs with the EGFP transgene were functionally titered in HeLa cells (VSVG-pseudotyped vectors) or 293T cells (BaEV-pseudotyped LVs). For LVs with other transgenes, a titer estimate was calculated based on the vectors’ p24-concentration related to the functional titer and p24 content of a control VSVG-pseudotyped vector (Table S3).

The FiCAR-T(R) lentiviral vector was produced at a GMP-competent contract development and manufacturing organization (CDMO; VIVEbiotech, Spain) and FiCAR-T_41BB by SIRION Biotech (part of Revvity; Germany).

### NK isolation and culturing

Natural killer (NK) cells were isolated from random healthy buffy coats using the Human NK Cell Isolation Kit (Miltenyi Biotec, Germany) and the manufacturer’s instructions. Subsequently, peripheral blood mononuclear cells (PBMCs) were obtained from one-day-old buffy coats provided by the Finnish Red Cross Blood Service under an institutional permit (FRCBS 178/6/2023). The PBMCs were separated using a Ficoll-based separation (Ficoll-Paque, Cytiva, Germany). After negative selection, the isolated NK cells were either directed for activation and transduction or cryopreserved in liquid nitrogen for further use.

The isolated NK cells were cultured and activated in an NK expansion medium consisting of NK MACS Basal Medium (Miltenyi Biotec, Germany) supplemented with 5% human AB serum (Sigma), 500 IU/ml IL-2 (proleukin S), 140 IU/ml IL-15 (Miltenyi Biotec, Germany), 100 U/ml penicillin and 100 ng/ml streptomycin (Penicillin-Streptomycin, Gibco, Thermo Fisher Scientific, Walthan, MA, USA).

### NK cell transduction

After three days of activation the NK cells were subjected to transduction using either the VSV-G- or BaEV-pseudotyped LVs. The transductions were performed in NK cell expansion medium, with a final cell density of 500,000 cells/ml, primarily using a multiplicity of infection (MOI) of 10. To enhance the transduction efficiency, we employed various enhancers and their combinations as follows:

Rosuvastatin (5 μM Cayman chemical, Ann Arbor, MI, USA) was added on day 0 (the day of NK cell isolation), and the cells activated in its presence until transduction on day 3. Before transduction, Retronectin (Takara, San Jose, Ca, USA) was coated onto the culture plates at a concentration of 25 μg/ml for two hours. Subsequently, the wells were blocked with 2% BSA (Gibco, Thermo Fisher Scientific, Walthan, MA, USA), and the plates used for transduction. Dextran (8 µg/ml, Sigma Aldrich, Merck, Germany) and the TLR4 inhibitors (1.1uM MRT67307, 5uM BX-795 or 24.8uM Amlexanox, all from MedChemExpress, Monmouth Junction, NJ, USA) were added 30 min before the transduction on day 3 and washed away on day 4. Vectofusin-1 (10 μg/ml, Miltenyi Biotec, Germany) was added simultaneously with the LVs on day 3. A subset of the NK cells was activated by co-culturing them with K562-based feeder cells (K562-mbIL-15-41BBL) in a 1:1 ratio from day 0 until the transduction on day 3. For the spinoculation, the NK cells were centrifuged at 400×g for 1–2 hours at RT after adding the LV.

Following transduction, the LVs and transduction enhancers were washed away using PBS at 24 hours after the transduction (day 4). Subsequently, the transduced cells were cocultured with feeder cells (K562-mbIL-15-41BBL) at a 1:2 NK:feeder ratio from day 4 until day 12. On day 12, an additional dose of feeder cells (also at a 1:2 ratio) was added. After two weeks of expansion, the CAR-NK cells were either analysed or cryopreserved. A subset of the CAR-NK cells was also analysed after one-week expansion following day 12.

### T cell isolation, culturing, and transduction

CAR-T cells were produced as previously described (Koski et al. 2025, submitted). Briefly, PBMCs were isolated from fresh buffy coats followed by T cell isolation by positive selection with CD4/CD8 microbeads (Miltenyi Biotec, Germany). Next, the isolated T cells were activated by incubating them for 24 hours with TransAct reagent (Miltenyi Biotec, Germany) at a density of 1×10^6 cells/ml in a complete T cell medium (TexMACS medium supplemented with 12 ng/ml of IL-7 and IL-15 (Miltenyi Biotec, Germany) and 100 U/ml of penicillin-streptomycin (Gibco, Thermo Fisher Scientific, Walthan, MA, USA)). The activated T cells were transduced using VSV-G-pseudotyped lentiviral vectors carrying the specific CARs. After 24 hours of transduction, the T cells were washed with medium without supplements and expanded in complete T cell medium for nine days prior to the analysis.

### Western blot

To analyse ASCT2 expression on NK cells, 2 x10^6^ cell samples were collected daily during NK cell activation to SDS sample buffer (Novagen, Thermo Fisher Scientific, Waltham, MA, USA), boiled for 10 minutes and stored for Western blotting. The Western blot analysis utilized an anti-ASCT2 antibody (1:1000, D7C12, 8057S, Cell Signalling Technology, Danvers, MA, USA) and anti-rabbit secondary antibody (1:800, 4412S, Cell Signalling Technology, Danvers, MA, USA).

### Flow cytometry analysis

The viability of the CAR-NK and -T cells was assessed by staining the cells with Zombie Green or Zombie Aqua (BioLegend, Germany) at a 1:600 dilution (10 minutes at room temperature, respectively). Following viability staining, the Fc receptors of the NK cells were blocked by incubating the cells with Fc Receptor Blocking Solution (Biolegend, Germany) at a dilution of approximately 1:30 for 10 minutes at room temperature. The actual antibody staining was performed for 15 minutes at room temperature or 30 minutes at +4°C in FACS buffer (PBS supplemented with 40 ng/ml EDTA and 5 mg/ml human albumin).

To detect lineage markers, the CAR-NK cells were stained with anti-CD56 (B339013, Biolegend, Germany), and CAR-T cells with anti-CD3 antibodies (300406, Biolegend, Germany). The CAR expression of both the CAR-NK and -T cells was assessed by staining the cells with an anti-FMC63 antibody (FM3-AY54P1, ACRO Biosystems, Newark, DE, USA). For phenotypic marker analysis, CAR-NK cells were stained with anti-CD16 (302048, Biolegend, Germany), -CD57 (393326, Biolegend, Germany), -LAG3 (369322, Biolegend, Germany), - NKG2A (130-113-567, Miltenyi Biotec, Germany), and -NKG2C (130-123-037, Miltenyi Biotec, Germany) antibodies.

The stained samples were analysed using the Symphony A1 instrument (BD Biosciences, Franklin Lakes, NJ, USA) with the relevant isotype, unstained, and untransduced control samples. The data analysis and compensations were performed using the FlowJo software (version 10, BD Biosciences, Franklin Lakes, NJ, USA) and isotype controls used for gating.

### Cytotoxicity assays

The cytotoxicity of the CAR-NK cells was evaluated using a luciferase-based assay. In brief, the luciferase expressing target cells (5000 RS4.11 or 20000 NALM6) were plated on a 384-well tissue culture plate in RPMI-1640 medium (Gibco, Thermo Fisher Scientific, Walthan, MA, USA) supplemented with 10% fetal bovine serum (Gibco, Thermo Fisher Scientific, Walthan, MA, USA) and 1% penicillin-streptomycin (Gibco, Thermo Fisher Scientific, Walthan, MA, USA). The CAR-NK cells were then added at an E:T ratio ranging from 0.12 to 10:1. After 4 hours of co-culture, the cells were lysed, and luciferase activity of the remaining live target cells measured following substrate addition (One-Glo, Promega, Madison, WI, USAS). The luminescence signal was quantified using the VICTOR Nivo Multimode Microplate Reader (PerkinElmer, Shelton, CT, USA). All measurements were performed in three technical replicates.

### Proteomic and Phosphoproteomic analyses

For the proteomic analyses, NK cells from three biological donors were transduced with the FiCAR-NK1 construct either without enhancers or using the BX795 and Retronectin combination, as previously described. Untransduced NK cells served as contros. On day 19, 10 x 10^6 CAR-NK cell pellets were collected in four technical replicates per condition, pelleted, snap-frozen in liquid nitrogen, and stored at −80°C until analysis.

### Mass spectrometry

Pellets were lysed in 8.0 M urea (#U5378-500G, Sigma Aldrich) in 100 mM ammonium biocarbonate (NH4HCO3, #A6141, Sigma Aldrich) with sonication (2 x 10min). Total protein concentration of the homogenates was measured with Bio-Rad Protein Assay Dye (#5000006, Bio-Rad Laboratories). The urea concentration was diluted to 1 M with 100 mM NH4HCO3. All the samples were reduced with 5 mM Tris(2-carboxyethyl)phosphine hydrochloride (#20490, Thermo Scientific), alkylated with 10 mM iodoacetamide (#122271000, Acros Organics) at room temperature, and trypsin-digested at 37°C for 16 hours using Sequencing Grade Modified Trypsin (V5113, Promega). After digestion, samples were acidified with 10% trifluoroacetic acid (TFA, #85049.051, VWR) and desalted with BioPureSPN MACRO TARGA C18 columns (HMM S18R, Nest Group) according to manufacturer’s instructions. From the eluate, one sixth was dried in a centrifuge concentrator (Concentrator Plus, Eppendorf) for total protein analysis. The rest was first enriched for phosphopeptides (The dried peptides were reconstituted in 30 µl buffer A (0.1% (vol/vol) TFA, 1% (vol/vol) acetonitrile (#83640.320, VWR) in HPLC grade water (#10505904, Fisher Scientific)). 50 µg of total protein from each sample was taken for total protein analysis, and 250 µg were used for phosphopeptide enrichment.

For phosphopeptide enrichment, the peptide mixture was loaded onto stage-tips packed with TiO_4_-IMAC microspheres, prewashed with loading buffer (80% acetonitrile/6% trifluoroacetic acid, TFA). TiO_4_-IMAC microspheres selectively captured phosphopeptides, while nonspecific peptides were removed with sequential washes of 50% acetonitrile/6% TFA/200 mM NaCl and 50% acetonitrile/0.1% TFA. Phosphopeptides were eluted using 10% NH_4_OH, followed by centrifugation to collect the eluate. The eluate was dried in a centrifuge concentrator (Concentrator Plus, Eppendorf) and stored for subsequent LC-MS/MS analysis.

Peptide samples were resuspended in buffer A (0.1 % FA, 1 % ACN, 30 µl for total, 60 µl for phospho). The samples were then diluted into a final concentration of 5 ng/µl and 20 µl was loaded into Evotip PURE (EV2011, Evosep) according to manufacturer instructions. Samples were then analyzed using the Evosep One liquid chromatography system coupled to a hybrid trapped ion mobility quadrupole TOF mass spectrometer (Bruker timsTOF Pro 2, Bruker Daltonics) (Meier, Brunner et al., 2018) via a CaptiveSpray nano-electrospray ion source (Bruker Daltonics). An 8⍰cm × 150⍰µm column with 1.5⍰µm C18 beads (EV1109, Evosep) was used for peptide separation with the 60 samples per day method for total samples and 30 samples per day method for phospho. Mobile phases A and B were 0.1 % formic acid in water and 0.1 % formic acid in acetonitrile, respectively. The MS analysis was performed in the positive-ion mode with dia-PASEF method (Meier, Brunner et al., 2020) with sample optimized data independent analysis (dia) scan parameters. To perform sample specific dia-PASEF parameter adjustment the default dia-short-gradient acquisition methods was adjusted based on the sample specific DDA-PASEF run with the software “tims Control” (Bruker Daltonics). The following parameters were modified for each sample type: m/z range; mass steps per cycle; mean cycle time (for total: da 400 – 1258, overlap 0.5, mass width 25; steps 13; cycle time 1.48 s. Phospho: da 345 – 1321, overlap 1.0, mass width 26; steps 22; cycle time 2.44 s. The ion mobility windows were set to best match the ion cloud density from the sample type specific DDA-runs.

To analyze diaPASEF data, the raw data (.d) were processed with DIA-NN v18.1 (Demichev et al., 2020; Demichev et al., 2022) utilizing spectral library generated from the UniProt human proteome (UP000005640, downloaded 2.4.2024 as a FASTA file, 20410 proteins). During library generation following settings were used for total samples: fixed modifications: carbamidomethyl (C); variable modifications: acetyl (protein N-term), oxidation (M); enzyme:Trypsin/P; maximun missed cleavages:1; mass accuracy fixed to 1.5e-05 (MS2) and 1.5e-05 (MS1); Fragment m/z set to 200-1800; peptide length set to 7-50; precursor m/z set to 300-1800; Precursor changes set to 1-4; protein inference not performed. For phospho data: fixed modifications: carbamidomethyl (C); variable modifications: acetyl (protein N-term),Phospho (STY); enzyme:Trypsin/P; maximun missed cleavages:1; mass accuracy fixed to 1.5e-05 (MS2) and 1.5e-05 (MS1); Fragment m/z set to 200-1800; peptide length set to 7-50; precursor m/z set to 300-1800; Precursor changes set to 2-4; protein inference not performed. All other settings were left to default.

### Vector copy number analysis

The absolute vector copy number in the cell population was determined using the QX200 digital droplet (dd)PCR system (Bio-Rad Laboratories, Hercules, CA, USA). One million cells were used to extract genomic DNA using the EZ1&2 DNA Tissue Kit for automated purification of DNA from cultured cells and the EZ2® Connect instrument according to manufacturer’s protocol (Qiagen, Germany). DeNovix DS-11 FX Spectrophotometer (DeNovix, Wilmington, USA) was used to determine the DNA concentration. The ddPCR assay was performed using the ddPCR Multiplex Supermix and probes detecting the vector sequence (HivPsi in FAM, assay ID: dEXD14812826) and housekeeping gene (RPP30 in HEX, assay ID: dHsaCP2500350) with the compatible restriction enzyme Hae III according to the manufacturer’s protocol (Bio-Rad Laboratories, Hercules, CA, USA). To generate droplets, the QX200 Automated droplet generator was used, and PCR amplification performed on the C1000 Touch Thermal cycler (Bio-Rad Laboratories, Hercules, CA, USA). The VCN was analysed using the Auto DG QX200 droplet reader PCR system and QuantaSoft software (Bio-Rad Laboratories, Hercules, CA, USA).

### Statistical analyses

Students t-test was used for comparison between groups. The level of statistical significance was set as p < 0.05. For proteomic analyses, protein-level intensity values were log2-transformed, and proteins that were not detected in at least 60% of samples of one sample group were discarded as low-quality identifications. Values were median normalized, and missing values were imputed using QRILC. Differential abundance analysis between sample groups was performed using a two-sample t-test, and p-values were adjusted using the Benjamini-Hochberg false discovery rate (FDR) procedure. Proteins were considered significantly differentially abundant if they met two criteria: an FDR-adjusted q-value below 0.01 and an absolute log2 fold change exceeding 1. Principal component analysis was used to visualise the differently abundant proteins. Functional enrichment was performed with DAVID.^27^

## Data availability statement

All essential data is provided within the manuscript and in supplemental material or can be obtained from the corresponding author upon reasonable request.

## Acknowledgments

This study was primarily conducted at Finnish Red Cross Blood Service, which we thank for the premises and financial support. The study was financially supported by the Finnish Cancer Foundation, Gilead foundation, Aamu Pediatric Cancer Foundation, the Foundation for Pediatric Research and Orion Corporation, which we warmly acknowledge. We thank Tiina Pohjankoski and Marita Kaislaranta for the research assistance and Anu Autio for her helpful comments during the project. The National Virus Vector Laboratory of the A.I.Virtanen Institute, University of Eastern Finland, Kuopio, Finland, and its former technician Anne Martikainen are acknowledged for lentiviral vector production and titering.

## Author contributions

HG, EJ, MK, KV, HP, DS, and SY designed the study. MK and KV supervised the study. EJ, FJ, MS, ME and HG designed and performed the experiments and analysed the data. EJ, FJ, MK and HG prepared the figures and wrote the manuscript. JK, MK, and HP designed the CAR constructs. AT, KS, and MV designed and performed the proteomics analyses, analysed its results and prepared related figures. DS coordinated and designed LVV production. MN performed the vector copy number analyses. LV contributed to data analysis. HC contributed to study design. All authors carefully revised the manuscript.

## Declaration of Generative AI and AI-assisted technologies in the writing process

During the preparation of this work the authors used Microsoft Copilot (2024-2025) to improve readability and language. After using this tool/service, the authors reviewed and edited the content as needed and take full responsibility for the content of the publication.

## References

1. Marin, D., Li, Y., Basar, R., Rafei, H., Daher, M., Dou, J., Mohanty, V., Dede, M., Nieto, Y., Uprety, N., et al. (2024). Safety, efficacy and determinants of response of allogeneic CD19-specific CAR-NK cells in CD19+ B cell tumors: a phase 1/2 trial. Nat Med 30, 772–784. 10.1038/s41591-023-02785-8.

2. Meeting highlights from the Pharmacovigilance Risk Assessment Committee (PRAC) 10-13 June 2024 | European Medicines Agency (EMA) https://www.ema.europa.eu/en/news/meeting-highlights-pharmacovigilance-risk-assessment-committee-prac-10-13-june-2024.

3. FDA Requires Boxed Warning for T cell Malignancies Following Treatment with BCMA-Directed or CD19-Directed Autologous Chimeric Antigen Receptor (CAR) T cell Immunotherapies | FDA https://www.fda.gov/vaccines-blood-biologics/safety-availability-biologics/fda-requires-boxed-warning-t-cell-malignancies-following-treatment-bcma-directed-or-cd19-directed.

4. Peng, L., Sferruzza, G., Yang, L., Zhou, L., and Chen, S. (2024). CAR-T and CAR-NK as cellular cancer immunotherapy for solid tumors. Preprint at Springer Nature, https://doi.org/10.1038/s41423-024-01207-0 10.1038/s41423-024-01207-0.

5. Gong, Y., Klein Wolterink, R.G.J., Wang, J., Bos, G.M.J., and Germeraad, W.T. V (2020). Chimeric antigen receptor natural killer (CAR-NK) cell design and engineering for cancer therapy. J Hematol Oncol 14, 73. 10.1186/s13045-021-01083-5.

6. Milone, M.C., and O’Doherty, U. (2018). Clinical use of lentiviral vectors. Leukemia 2018 32:7 32, 1529–1541. 10.1038/s41375-018-0106-0.

7. Gutierrez-Guerrero, A., Cosset, F.L., and Verhoeyen, E. (2020). Lentiviral Vector Pseudotypes: Precious Tools to Improve Gene Modification of Hematopoietic Cells for Research and Gene Therapy. Viruses 2020, Vol. 12, Page 1016 12, 1016. 10.3390/V12091016.

8. Deng, L., Liang, P., and Cui, H. (2023). Pseudotyped lentiviral vectors: Ready for translation into targeted cancer gene therapy? Genes Dis 10, 1937–1955. 10.1016/J.GENDIS.2022.03.007.

9. Finkelshtein, D., Werman, A., Novick, D., Barak, S., and Rubinstein, M. (2013). LDL receptor and its family members serve as the cellular receptors for vesicular stomatitis virus. Proc Natl Acad Sci U S A 110, 7306–7311. 10.1073/pnas.1214441110.

10. Gong, Y., Klein Wolterink, R.G.J., Janssen, I., Groot, A.J., Bos, G.M.J., and Germeraad, W.T.V. (2020). Rosuvastatin Enhances VSV-G Lentiviral Transduction of NK Cells via Upregulation of the Low-Density Lipoprotein Receptor. Mol Ther Methods Clin Dev 17, 634. 10.1016/J.OMTM.2020.03.017.

11. Bari, R., Granzin, M., Tsang, K.S., Roy, A., Krueger, W., Orentas, R., Pfeifer, R., Moeker, N., Verhoeyen, E., Dropulic, B., et al. (2019). A distinct subset of highly proliferative and lentiviral vector (LV)-transducible NK cells define a readily engineered subset for adoptive cellular therapy. Front Immunol 10, 471032. 10.3389/FIMMU.2019.02001/BIBTEX.

12. Allan, D.S.J., Chakraborty, M., Waller, G.C., Hochman, M.J., Poolcharoen, A., Reger, R.N., and Childs, R.W. (2021). Systematic improvements in lentiviral transduction of primary human natural killer cells undergoing ex vivo expansion. Mol Ther Methods Clin Dev 20, 559. 10.1016/J.OMTM.2021.01.008.

13. Chockley, P., Patil, S.L., and Gottschalk, S. (2021). Transient blockade of TBK1/IKK epsilon allows efficient transduction of primary human NK cells with VSV-G-pseudotyped lentiviral vectors. Cytotherapy 23, 787. 10.1016/J.JCYT.2021.04.010.

14. Hanenberg, H., Xiao, X.L., Dilloo, D., Hashino, K., Kato, I., and Williams, D.A. (1996). Colocalization of retrovirus and target cells on specific fibronectin fragments increases genetic transduction of mammalian cells. Nature Medicine 1996 2:8 2, 876–882. 10.1038/nm0896-876.

15. Fenard, D., Ingrao, D., Seye, A., Buisset, J., Genries, S., Martin, S., Kichler, A., and Galy, A. (2013). Vectofusin-1, a New Viral Entry Enhancer, Strongly Promotes Lentiviral Transduction of Human Hematopoietic Stem Cells. Mol Ther Nucleic Acids 2, e90. 10.1038/MTNA.2013.17.

16. Koski, J., Jahan, F., Luostarinen, A., Schenkwein, D., Ylä-Herttuala, S., Göös, H., Monzo, H., Ojala, P.M., Maliniemi, P., and Korhonen, M. (2022). Novel modular chimeric antigen receptor spacer for T cells derived from signal regulatory protein alpha Ig-like domains. Frontiers in Molecular Medicine 2. 10.3389/fmmed.2022.1049580.

17. Jahan, F., Koski, J., Schenkwein, D., Ylä-Herttuala, S., Göös, H., Huuskonen, S., Varjosalo, M., Maliniemi, P., Leitner, J., Steinberger, P., et al. (2023). Using the Jurkat reporter T cell line for evaluating the functionality of novel chimeric antigen receptors. Frontiers in Molecular Medicine 3. 10.3389/FMMED.2023.1070384.

18. Colamartino, A.B.L., Lemieux, W., Bifsha, P., Nicoletti, S., Chakravarti, N., Sanz, J., Roméro, H., Selleri, S., Béland, K., Guiot, M., et al. (2019). Efficient and Robust NK-Cell Transduction With Baboon Envelope Pseudotyped Lentivector. Front Immunol 10, 2873. 10.3389/FIMMU.2019.02873.

19. Schwarzwaelder, K., Howe, S.J., Schmidt, M., Brugman, M.H., Deichmann, A., Glimm, H., Schmidt, S., Prinz, C., Wissler, M., King, D.J.S., et al. (2007). Gammaretrovirus-mediated correction of SCID-X1 is associated with skewed vector integration site distribution in vivo. Journal of Clinical Investigation 117, 2241. 10.1172/JCI31661.

20. Liu, E., Tong, Y., Dotti, G., Shaim, H., Savoldo, B., Mukherjee, M., Orange, J., Wan, X., Lu, X., Reynolds, A., et al. (2018). Cord blood NK cells engineered to express IL-15 and a CD19-targeted CAR show long-term persistence and potent anti-tumor activity. Leukemia 32, 520. 10.1038/LEU.2017.226.

21. Müller, S., Bexte, T., Gebel, V., Kalensee, F., Stolzenberg, E., Hartmann, J., Koehl, U., Schambach, A., Wels, W.S., Modlich, U., et al. (2020). High Cytotoxic Efficiency of Lentivirally and Alpharetrovirally Engineered CD19-Specific Chimeric Antigen Receptor Natural Killer Cells Against Acute Lymphoblastic Leukemia. Front Immunol 10, 3123. 10.3389/FIMMU.2019.03123/FULL.

22. Albinger, N., Müller, S., Kostyra, J., Kuska, J., Mertlitz, S., Penack, O., Zhang, C., Möker, N., and Ullrich, E. (2024). Manufacturing of primary CAR-NK cells in an automated system for the treatment of acute myeloid leukemia. Bone Marrow Transplant 59, 489. 10.1038/S41409-023-02180-4.

23. Sutlu, T., Nyström, S., Gilljam, M., Stellan, B., Applequist, S.E., and Alici, E. (2012). Inhibition of Intracellular Antiviral Defense Mechanisms Augments Lentiviral Transduction of Human Natural Killer Cells: Implications for Gene Therapy. Hum Gene Ther 23, 1090. 10.1089/HUM.2012.080.

24. Heron, L., Virsolvy, A., Peyrollier, K., Gribble, F.M., Cam, A. Le, Ashcroft, F.M., and Bataille, D. (1998). Human α-endosulfine, a possible regulator of sulfonylurea-sensitive KATP channel: Molecular cloning, expression and biological properties. Proc Natl Acad Sci U S A 95, 8387. 10.1073/PNAS.95.14.8387.

25. Imai, C., Iwamoto, S., and Campana, D. (2005). Genetic modification of primary natural killer cells overcomes inhibitory signals and induces specific killing of leukemic cells. Blood 106, 376–383. 10.1182/blood-2004-12-4797.

26. Follenzi, A., and Naldini, L. (2002). [26] Generation of HIV-1 derived lentiviral vectors. Methods Enzymol 346, 454–465. 10.1016/S0076-6879(02)46071-5.

27. Sherman, B.T., Hao, M., Qiu, J., Jiao, X., Baseler, M.W., Lane, H.C., Imamichi, T., and Chang, W. (2022). DAVID: a web server for functional enrichment analysis and functional annotation of gene lists (2021 update). Nucleic Acids Res 50, W216–W221. 10.1093/NAR/GKAC194,.

