## Supplemental material for "Efficient NK cell transduction with VSV-G-pseudotyped lentiviral vectors"

**Table S1. Total protein analysis of differentially transduced NK cells**

A) Proteins identied in total protein lysate analysis. Means and standard deviations of normalised label free quantified (LFQ) values (log2 transformed) from four technical replicated from three donors. Each condition (UTD= untransduced, woE= without enhancer transduced and BX+R= with BX795 and Retronectin transduced cells) is shown in separate columns. NA values imputed as described in material and methods.

B) Comparison of protein expression between untransduced and without enhancers transduced NK cells. Fold changes (log2), p-values, sample mean values and control mean values are shown as separete columns.

C) Comparison of protein expression between without enhancers and BX795/Retronectin transduced NK cells. Fold changes (log2), p-values, sample mean values and control mean values are shown as separete columns.

D) Enriched Bioogical Process Gene Ontology (BP-GO) terms from total protein analysis. Analysis was performed using The Database for Annotation, Visualization and Integrated Discovery (DAVID) (https://david.ncifcrf.gov/).

E) Enriched KEGG-pathways from total protein analysis. Analysis was performed using The Database for Annotation, Visualization and Integrated Discovery (DAVID) (https://david.ncifcrf.gov/).

F) Enriched REACTOME-pathways from total protein analysis. Analysis was performed using The Database for Annotation, Visualization and Integrated Discovery (DAVID) (https://david.ncifcrf.gov/).

**Table S2. Phosphoprotein analysis of differentially transduced NK cells.**

A) Proteins identied in phosphoprotein analysis. Means and standard deviations of normalised MS1 quantified values  (log2 transformed) ) from four technical replicated from three donors. Each condition (UTD= untransduced, woE= without enhancer transduced and BX+R= with BX795 and Retronectin transduced cells) is shown in separate columns. NA values imputed as described in material and methods.

B) Comparison of phosphoproteins between untransduced and without enhancers transduced NK cells. Fold changes (log2), p-values, sample mean values and control mean values are shown as separete columns.

C) Comparison of phosphoproteins between without enhancers and BX795/retronectin transduced NK cells. Fold changes (log2), p-values, sample mean values and control mean values are shown as separete columns.

**Table S3. Functional titers (TU/ml) and estimated functional titers of lentivirus vectors used in this study.** TU = transduction units.

| **Vector group** | **Name** | **Functional titer (TU/ml)** | **Titer estimate (TU/ml)** |
|---|---|---|---|
| VSVG.LVs | VSVG-GFP | 2,70E+09 | 1,3E+09 |
| | VSV-G MOCK | | 2,20E+09 |
| | VSV_G NK1 | | 2,00E+09 |
| | VSV_G NK2 | | 1,30E+09 |
| | VSV_G NK3 | | 2,40E+09 |
| | VSV_G FiCAR 1v3 | | 1,10E+09 |
| BaEV.LVs; Production 1 | BaEVRless-GFP | 7,16E+06 | 3,35E+08 |
| | BaEVRlessMOCK | | 4,95E+08 |
| | BaEVRlessNK1 | | 2,53E+08 |
| | BaEVRlessNK2 | | 2,23E+08 |
| | BaEVRlessNK3 | | 8,33E+08 |
| | BaEVRlessFICAR1V3 | | 1,86E+08 |
| BaEV.LVs; Production 2 | BaEVRless-GFP | 1,10E+06 | 6,70E+07 |
| | BaEVRlessMOCK | | 7,30E+08 |
| | BaEVRlessNK1 | | 3,60E+08 |
| | BaEVRlessNK2 | | 3,20E+07 |
| | BaEVRlessNK3 | | 8,10E+07 |
| | BaEVRlessFICAR1V3 | | 3,20E+07 |

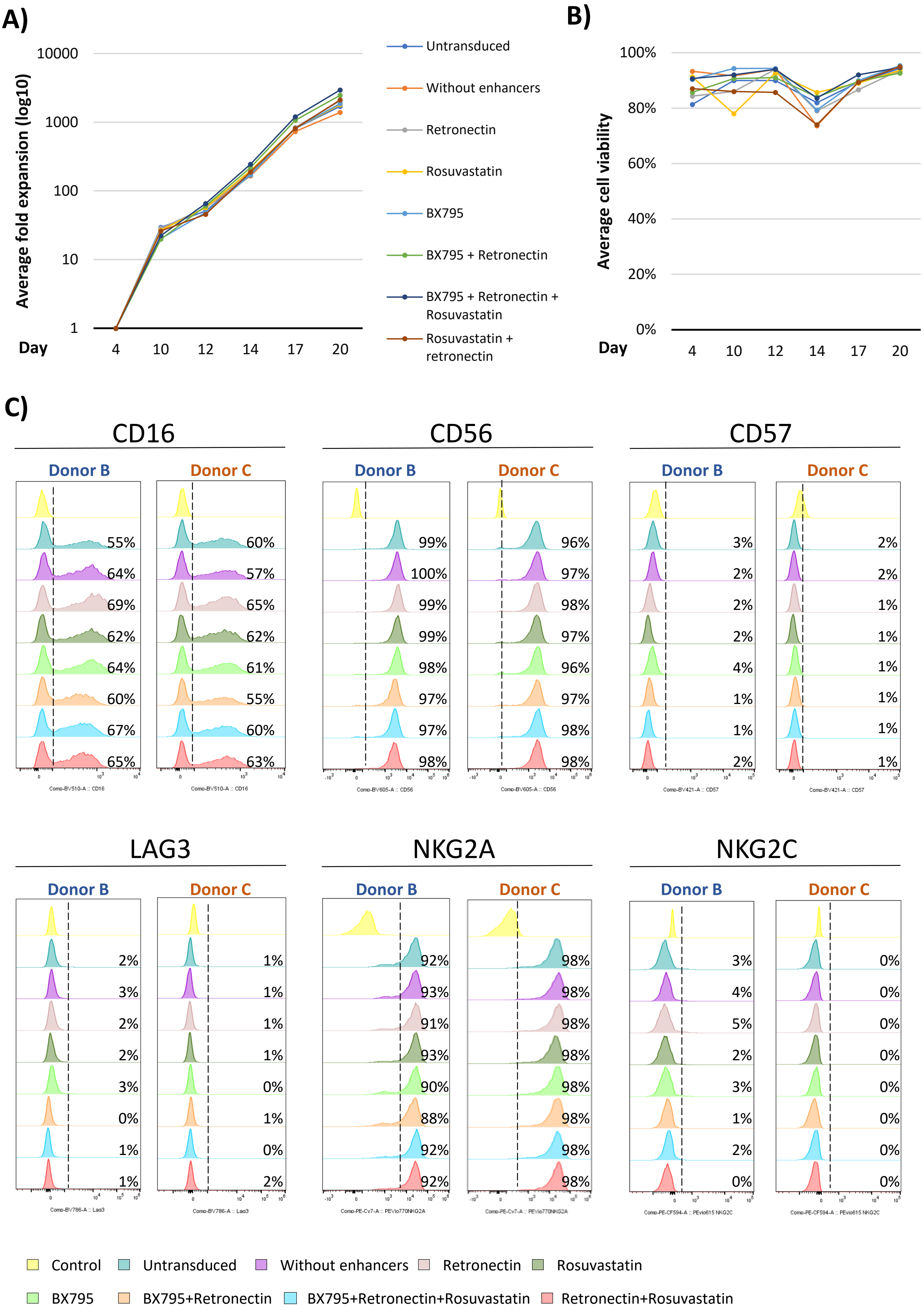

A)
Average fold expansion (log10)
10000
1000
100
10
1
Day
4
10
12
14
17
20
Untransduced
Without enhancers
Retronectin
Rosuvastatin
BX795
BX795 + Retronectin
BX795 + Retronectin + Rosuvastatin
Rosuvastatin + retronectin
B)
Average cell viability
100%
80%
60%
40%
20%
0%
Day
4
10
12
14
17
20
C)
CD16
Donor B
55% 64% 69% 62% 64% 60% 67% 65%
Donor C
60% 57% 65% 62% 61% 55% 60% 63%
CD56
Donor B
99% 100% 99% 99% 98% 97% 97% 98%
Donor C
96% 97% 98% 97% 96% 97% 98% 98%
CD57
Donor B
3% 2% 2% 2% 4% 1% 1% 2%
Donor C
2% 2% 1% 1% 1% 1% 1% 1%
LAG3
Donor B
2% 3% 2% 2% 3% 0% 1% 1%
Donor C
1% 1% 1% 1% 0% 1% 0% 2%
NKG2A
Donor B
92% 93% 91% 93% 90% 88% 92% 92%
Donor C
98% 98% 98% 98% 98% 98% 98% 98%
NKG2C
Donor B
3% 4% 5% 2% 3% 1% 2% 0%
Donor C
0% 0% 0% 0% 0% 0% 0% 0%
Control
Untransduced
Without enhancers
Retronectin
Rosuvastatin
BX795
BX795+Retronectin
BX795+Retronectin+Rosuvastatin
Retronectin+Rosuvastatin

**Figure S1. NK cells transduced with CAR**

A) Fold expansion of after transduction of NK cells during days 4-20. Feeder cells were added at 1:2 on days 4 and 12. Cumulative fold expansions for each transduction method are shown with color-coded lines (keys in the panel on the right). Colored dots show the average expansion of cells derived from two donors.

B) Average viability of CAR-NK cells transduced using different transduction enhancers. Cells were stained with Trypan Blue and counted. Percentage of viable cells (average of two biological replicates) is shown on the Y-axis and viability curves for individual transduction method are color coded (keys in the panel on the right).

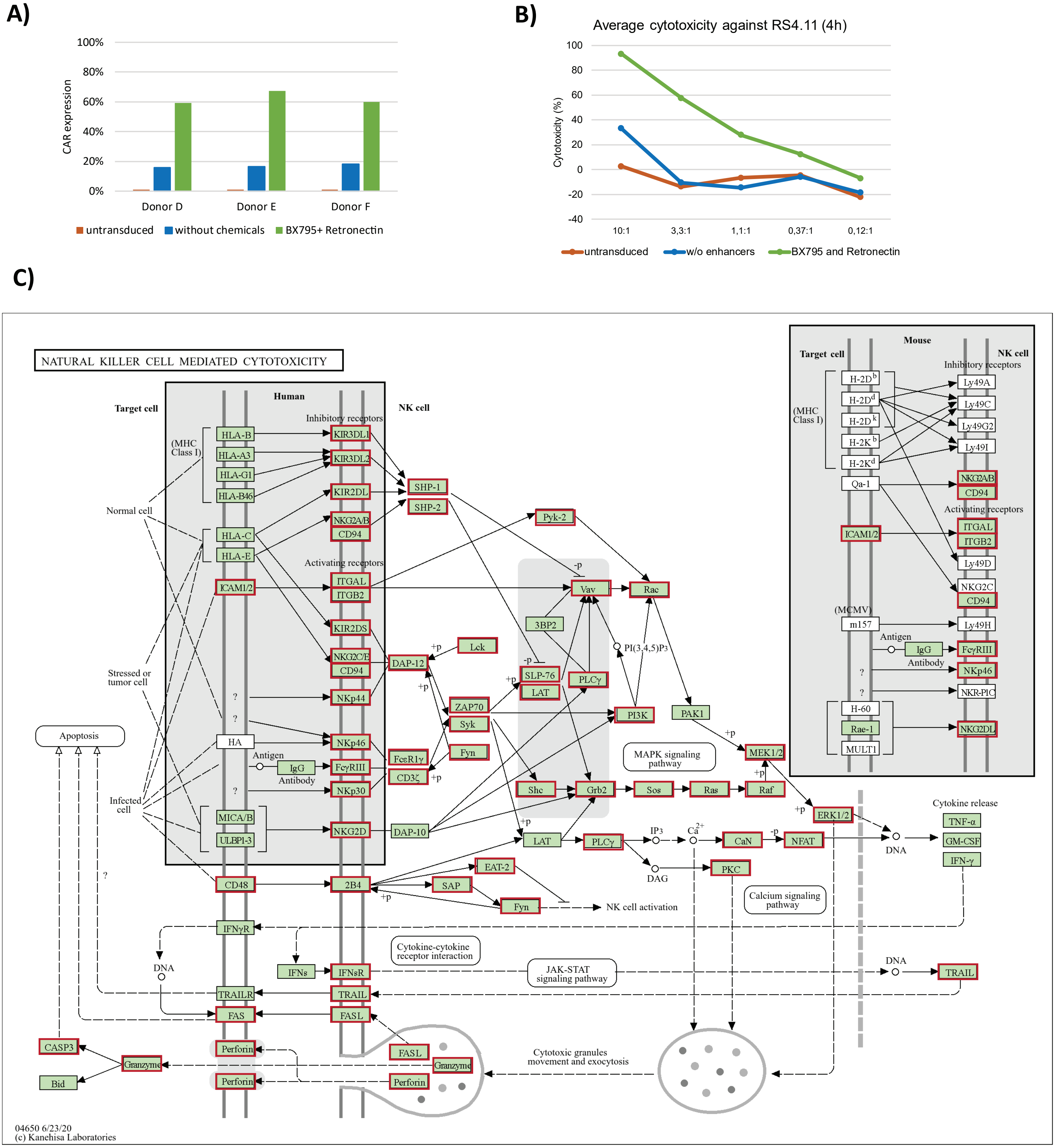


Downloaded : https://www.genome.jp/pathway/hsa04650

**Figure S2. Proteomic profiling of CAR-NK cells transduced with BX795 and Retronectin**

A) CAR expression and B) cytotoxicity of CAR-NK cells used in experiment. Average of three biological donors (each was performed using three technical replicates) C) KEGG-pathway map of natural killer cell mediated cytotoxicity. Proteins identified in total protein lysate analysis are highlighted with red boxes. Map downloaded from https://www.genome.jp/pathway/hsa04650.

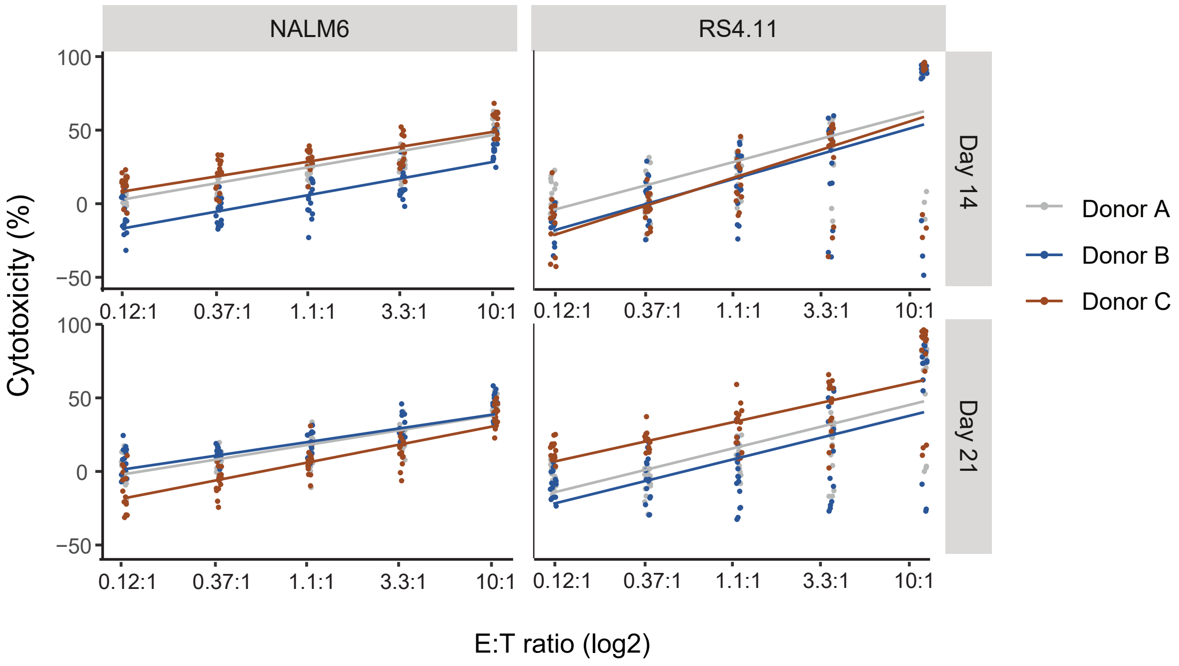


**Figure S3. Cytotoxicity of d14 and d20 CAR-NK cells from different donors against RS4.11 and NALM6 ALL cell lines.**

CAR-NK cells with different constructs were incubated with target cells for four hours. For each donor (n = 3, color-coded), linear regression was performed across five construct groups, each with three technical replicates. Cytotoxicity was assayed on day 14 (upper graphs) and 21 (lower graphs). Effector to target ratios (E:T) in log2 scale are shown on the X-axis.

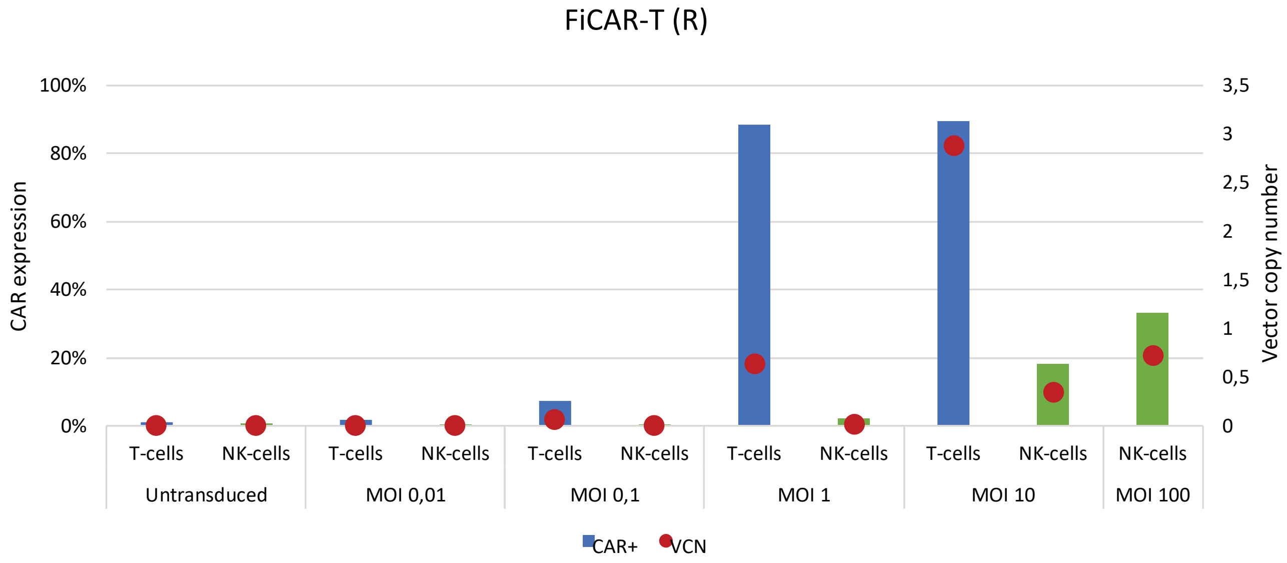


**Figure S4. FiCAR-T(R) titration using NK and T cells.**

NK and T cell CAR expression after titration with FiCAR-T(R) -LV. following IL-2/IL-15 activation NK cells were transduced in the presence of of BX795 and Retronectin. T cells were activated with transact-beads prior to transduction. CAR expression was assessed on day 12 for NK cells and on day 9 for T cells by flow cytometry. N=1

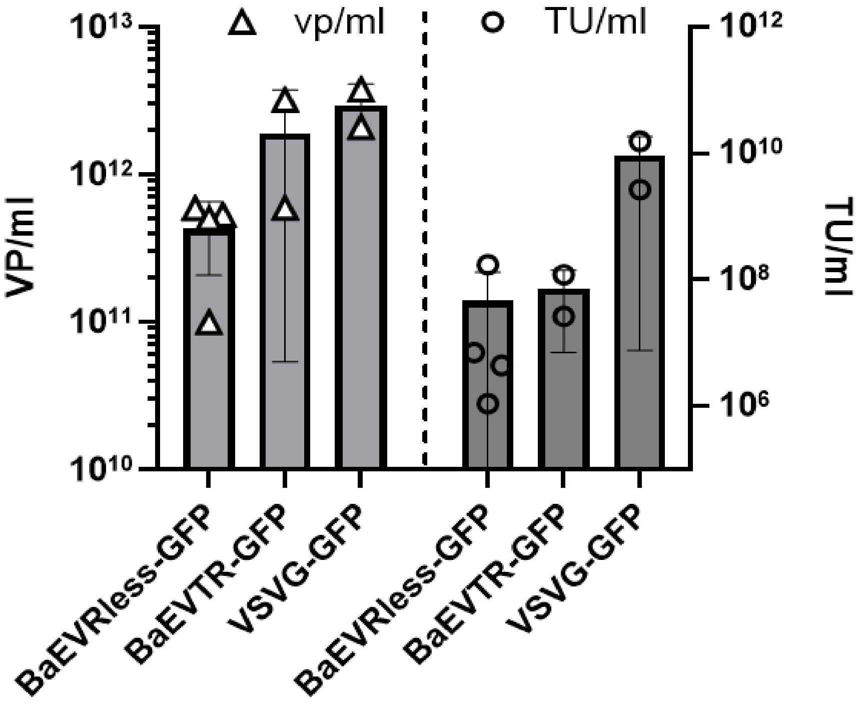


**Figure S5. Analytics of produced BaEV-LV lots.**

Vector particles (vp) per milliliter was calculated based on the p24 concentration of the vector preparations. Functional titers expressed as transducing units per milliliter (TU/ml) were determined in transduced cells.

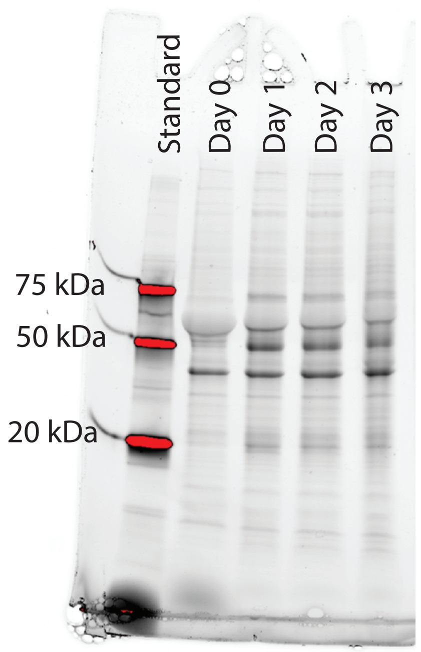


**Figure S6. SDS-PAGE gel loading control.**

Stain-free SDS-PAGE gel imaged directly after the gel running shows equal sample loading between the samples.
